# Causal roles of prefrontal cortex during spontaneous perceptual switching are determined by brain state dynamics

**DOI:** 10.1101/2021.04.16.440188

**Authors:** Takamitsu Watanabe

## Abstract

The prefrontal cortex (PFC) is thought to orchestrate cognitive dynamics. However, in tests of bistable visual perception, no direct evidence supporting such presumable causal roles of the PFC has been reported. Here, using a novel brain-state-dependent neural stimulation system, we found that three PFC regions—right frontal eye fields and anterior/posterior dorsolateral PFCs (a/pDLPFCs)—have causal effects on perceptual dynamics but the behavioural effects are detectable only when we modulated the PFC activity in brain-state-/state-history-dependent manners. Also, we revealed that the brain-dynamics-dependent behavioural causality is underpinned by transient changes in the brain state dynamics, and such neural changes are determined by structural transformations of hypothetical energy landscapes. Moreover, we identified different functions of the three PFC areas: in particular, we found that aDLPFC enhances the integration of the two PFC-active brain states, whereas pDLPFC promotes the diversity between them. This work resolves the controversy over the PFC roles in spontaneous perceptual switching and underlines brain state dynamics in fine investigations of brain-behaviour causality.

**Impact statement:** Prefrontal causal roles are changing during bistable visual perception, which was determined by large-scale brain state dynamics and attributable to hypothetical energy landscapes that underpin the brain state dynamics.

## Introduction

Dynamic and flexible changes are among fundamental key properties of human cognition and perception. Bistable visual perception has been widely used to investigate such cognitive dynamics ^1,2^, and the prefrontal cortex (PFC)—in particular, right frontal eye fields (FEF) and anterior/posterior dorsolateral PFC (a/pDLPFC)—has been thought to be involved in the spontaneous perceptual switching ^1–10^. Theoretical work also indicates that top-down signals from these PFC regions to the visual cortex are essential to the perceptual inference ^11,12^. These studies indicate that inhibitory neural modulation of the PFC areas should induce behavioural changes in the bistable visual perception.

However, no empirical human study has identified such behavioural causality of the PFC ^2^. Instead, research using transcranial magnetic stimulation (TMS) reported that neural suppression of a brain area near the right aDLPFC did not affect bistable visual perception ^13^, and other studies claimed that the PFC activity is not essential to the emergence of multistable perception ^14,15^ but mere a consequence of it ^15–18^.

Why is it so difficult to detect the prefrontal causal roles in the multistable perception paradigm? Here, given that the whole-brain neural activity during bistable perception is described as large-scale brain state dynamics ^19^, we hypothesise that causal roles of the PFC regions should also be dynamically changing during the fluctuating visual awareness. That is, if the neural activity in the multistable perception can be stated as dwelling in and transitions between a parsimonious number of brain states ^19^, the detectability of the prefrontal causal effects on the perceptual awareness should be determined by the brain state in which the neural activity pattern is staying when the neural stimulation is administered. If so, such state-dependent behavioural causality should be hardly observed when we intervene in the PFC activity without tracking the large-scale brain state dynamics.

We directly tested this hypothesis with a brain-state-dependent neural stimulation system ^20–22^. The system was devised through linking energy landscape analysis ^19,23–26^, which enabled us to identify and track the brain state dynamics during seemingly unstable behaviours, to an electroencephalogram (EEG)-triggered TMS ^27–30^, which allowed brain-activity-dependent non-invasive neural modulation.

To focus on neural mechanisms in the higher-order cortex, this study did not adopt a test of binocular rivalry, which has been often explained by inter-hemispheric neural suppression in the lower-level brain systems ^31–33^; instead, we used a test of bistable visual perception induced by a structure-from-motion (SFM) stimulus (Fig. 1a), in which the same visual stimulus is presented to both of the eyes of human participants.

**Fig. 1.**
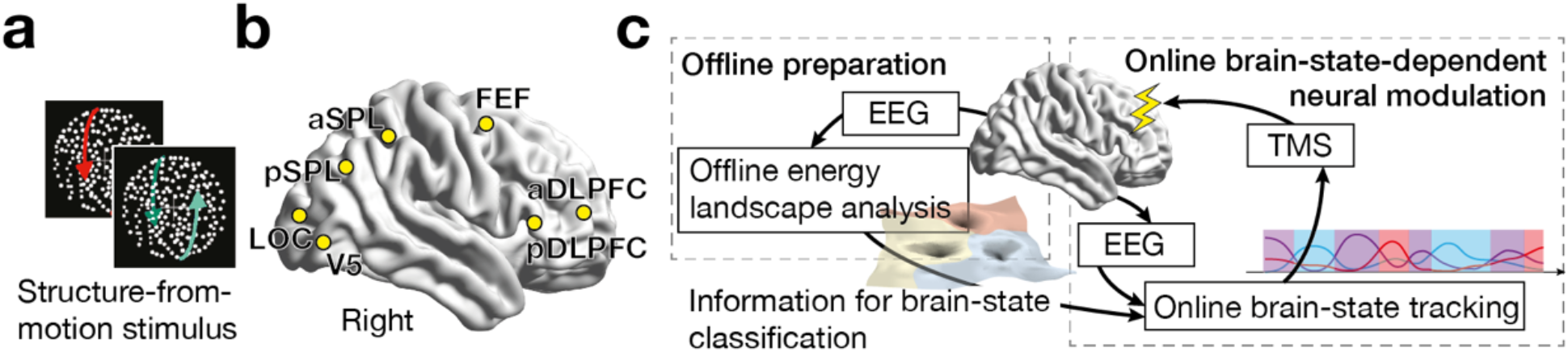
**a-b.** While the participants were presented with a structure-from-motion (SFM) stimulus (**panel a**), we recorded EEG signals from the seven brain regions (**panel b**). aDLPFC/pDLPFC, anterior/posterior dorsolateral prefrontal cortex; FEF, frontal eye fields; aSPL/pSPL, anterior/posterior superior parietal lobule; LOC, lateral occipital complex. **c.** We first conducted offline energy landscape analysis, whose results allowed us to categorise brain activity patterns into either of the major brain states. By implementing such classification information to an online EEG analysis, we tracked brain state dynamics and administered inhibitory TMS in state-/state-history-dependent manners.

## Results

### Individual brain state dynamics

As a preparation, we first conducted an offline analysis of EEG data recorded from the seven cortical regions ^19^(Fig. 1b) of 65 healthy adults while they were experiencing the SFM-induced bistable visual perception (left panel of Fig. 1c; Control Experiment I). The energy landscape analysis confirmed that the neural dynamics during the bistable perception were described as dwelling in and transitions between the following three major brain states (model fitting accuracy ≥ 84.1%; Figs. 1d and 1e): Frontal-area-dominant state (F state, 32.9±0.35% of all the brain states, mean±sem), Visual-area-dominant state (V state, 33.9±0.42%) and Intermediate state (Int state, 34.0±0.43%). The whole-brain neural activity pattern rarely moved between F and V state directly (0.8±0.007% of all the transitions). Instead, it almost always used Int state as a relay point (Figs. 1f and 1g). Moreover, the length of such an indirect neural travel between F and V state via Int state accurately predicted the percept duration (*r*_64_ = 0.67, *P* < 10^−5^; Fig. 1h).

**Fig. 1.**
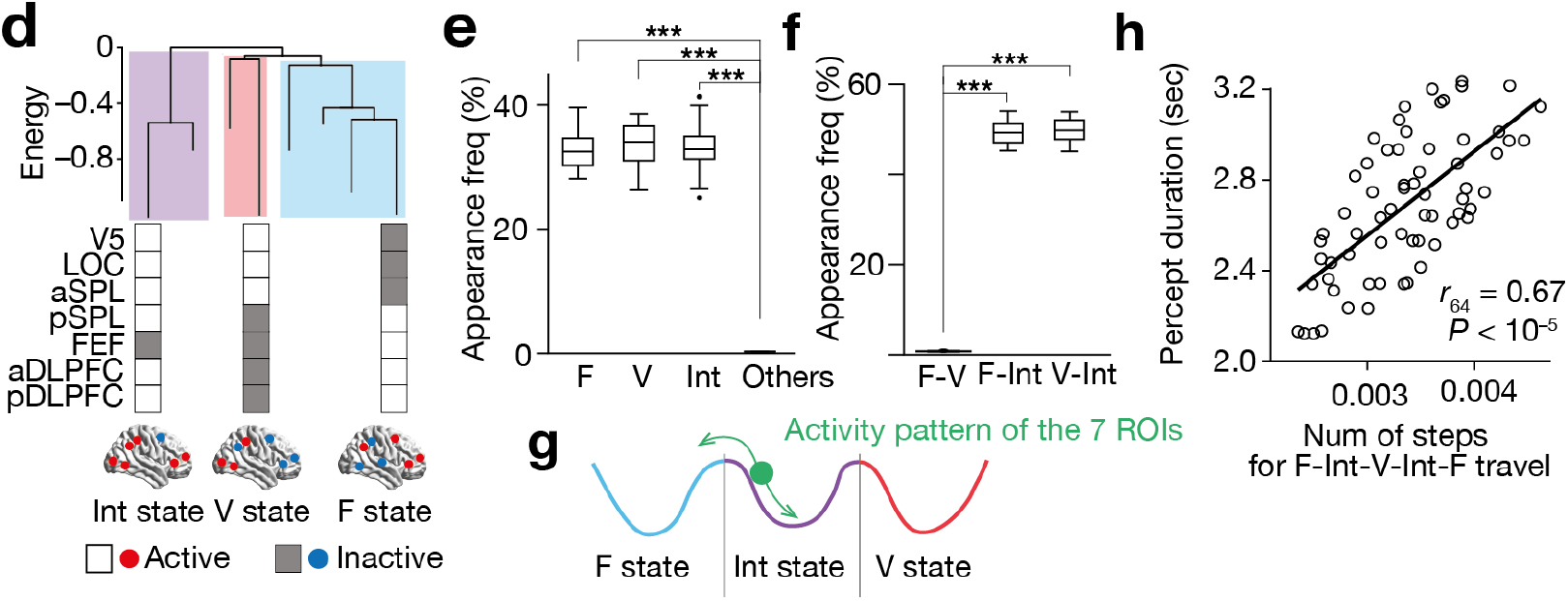
**d.** In Control Experiment I, we identified individual disconnectivity graph that determined the structure of the individual energy landscape (**panel d** for Participant #1). In the graph, a branch end represents a local minimum, and the lower energy values of the local minima indicate the more stable brain states; the heights of the energy barriers between the local minima are known to be inversely correlated with the transition frequencies between the brain states. **e-h.** Almost all the brain activity patterns were categorised into either of F, V or Int state, and the appearance frequencies of the other states were negligible (<0.1%) compared to the three major states (**panel e**). The direct transition between F and V state was rare (<0.9%; **panel f**). These results indicate that the brain activity pattern is travelling between F and V state via Int state (**panel g**). A random-walk simulation shows that the number of steps for such F-Int-V travel indicates the individual perceptual stability (**panel h**; each circle represents each participant). *** indicates *P*_Bonferroni_ < 0.001 in paired *t*-tests (*df*=64).

These EEG observations are essentially the same as our previous fMRI findings ^19^, which also provides face validation to the locations of the EEG electrodes.

### State-dependent neural stimulation

These individual energy landscape structures had sufficient information about how to categorise each neural activity pattern at each timepoint into either of the three major brain states. In Control Experiment II, we implemented such classification information into an online EEG analysis ^29,30^(right panel of Fig. 1c) and tracked the brain state dynamics with high accuracy (similarity to results based on the offline analysis >82.3%; Fig. 1i). Then, by linking the online analysis to monophasic TMS, we succussed in triggering a burst of inhibitory TMS only when the neural activity pattern was staying in a specific major brain state (accuracy >84.8%, Fig. 1j; latency <0.8msec, Fig. 1k).

**Fig. 1.**
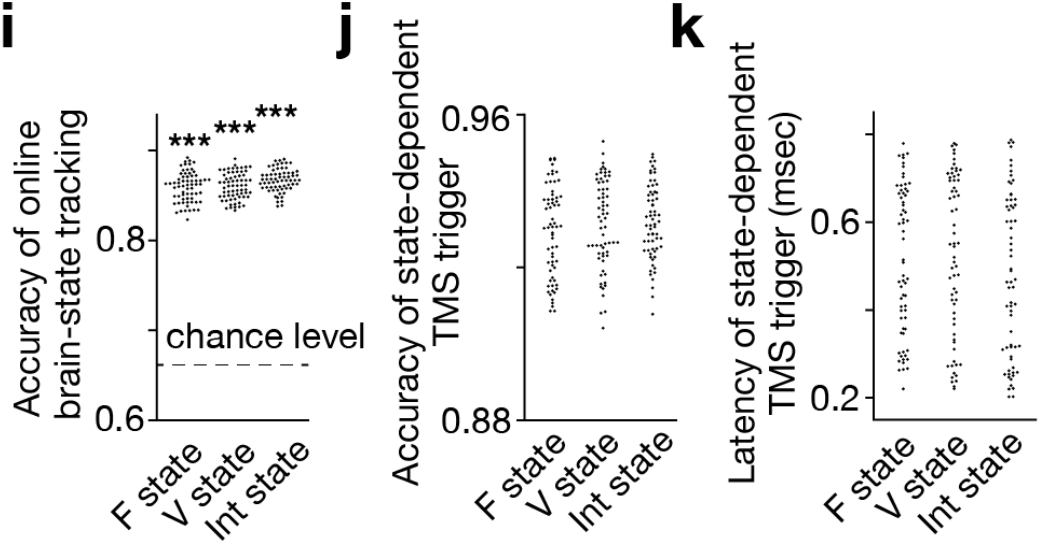
**i-k.** In Control Experiment II, the online EEG analysis could track brain state dynamics as accurately as the offline analysis (**panel i**). Brain-state-dependent TMS triggering system also achieved accurate neural tracking (**panel j**) with short delays (**panel k)**. Note that the accuracy in the TMS trigger (**panel j**) tended to be higher than that in simple brain-state tracking (**panel i**) presumably because TMS was triggered only when a specific brain state was expected to continue in a certain period (here, 150msec) and such a criterion should work as temporal smoothing and improve the signal-to-noise ratio. Each dot represents each participant. *** indicates *P*_Bonferroni_ < 0.001 in paired *t*-tests (*df* = 64).

### State-dependent causality

By applying this state-dependent TMS over the three PFC regions (i.e., aDLPFC, pDLPFC and FEF; Fig. 1l), we found that the three prefrontal areas had causal behavioural effects on the spontaneous perceptual switching in a brain-state-dependent manner (*N*=34; *F*_8,33_=25.9, *P*<10^−3^ for the main effect in a two-way ANOVA; Fig. 2a).

**Fig. 1.**
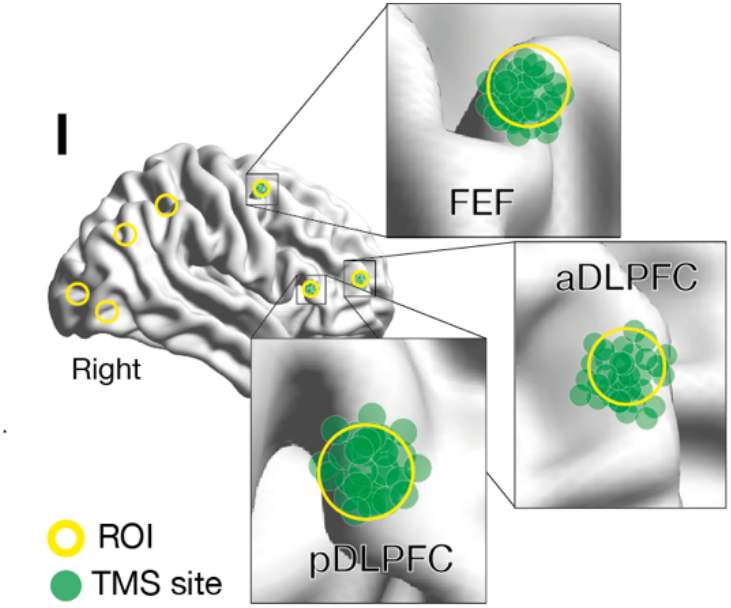
**l.** We administered inhibitory TMS over the three PFC regions. In one TMS condition, we placed the TMS coil over one of the PFC areas using a stereoscopic neuro-navigation system. Each green circle represents the coordinates of the stimulated brain site for an individual participant, which was re-measured with the neuro-navigation system at the end of each experiment day and averaged across the four-day sessions in the main experiment. The green circles were mapped closely onto the original coordinates (the centres of the yellow circles).

**Fig. 2.**
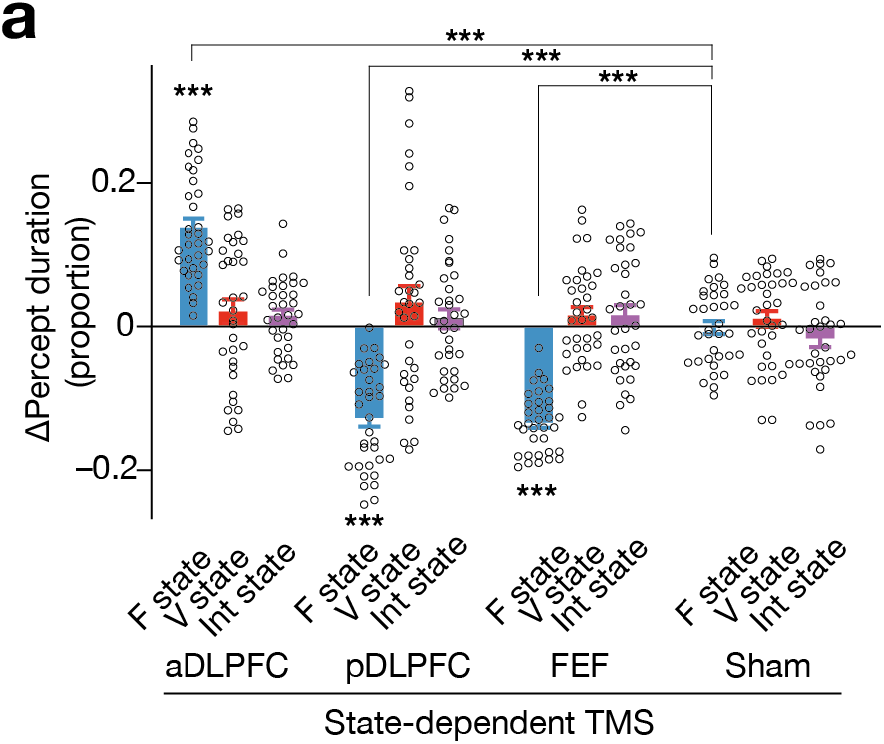
**a.** State-dependent TMS revealed that causal behavioural roles of the three PFC regions are detectable when the brain activity pattern stays in F state. The Y-axis shows the proportional changes in the median percept duration (= (TMS–control)/control). The X-axis indicates the TMS conditions. *** indicates *P*_Bonferroni_ < 0.001 in paired *t*-tests (*df*=33). Each circle represents each participant. The error bars show the s.e.m.

The neural inhibition of aDLPFC during F state prolonged the percept duration (*t*_33_=9.4, *P*_Bonferroni_<0.001 in a post-hoc paired *t*-test, Cohen’s *d*=1.7), whereas that during V or Int state induced no behavioural change (*t*_33_<1.7, *P*_Bonferroni_>0.05, *d*<0.3).

Regarding pDLPFC, the F-state-dependent TMS destabilised the visual perception (*t*_33_=10.4, *P*_Bonferroni_<0.001, *d*=1.8), whilst neither V- or Int-state-dependent TMS induced any behavioural effect (*t*_33_<1.3, *P*_Bonferroni_>0.05, *d*<0.29).

The F-state-dependent TMS over FEF reduced the percept duration (*t*_33_=10.5, *P*_Bonferroni_<0.001, *d*=1.9), whereas no behavioural change was observed in the other FEF TMS conditions (*t*_33_<0.8, *P*_Bonferroni_>0.05, *d*<0.39)

### State-history-dependent causality

If some behavioural causalities are detectable when the whole-brain neural activity pattern is dwelling in specific brain states, others may emerge when the neural activity pattern has finished travelling a particular brain state trajectory. We then tested this hypothesis and found such state-history-dependent causality (*F*_5,33_=85.6, *P*<10^−3^ for the main effect in a two-way ANOVA; Fig. 2b). Here, we focused on Int state because only the brain state had two incoming pathways (i.e., a path from F state and one from V state; Fig. 1g).

**Fig. 2.**
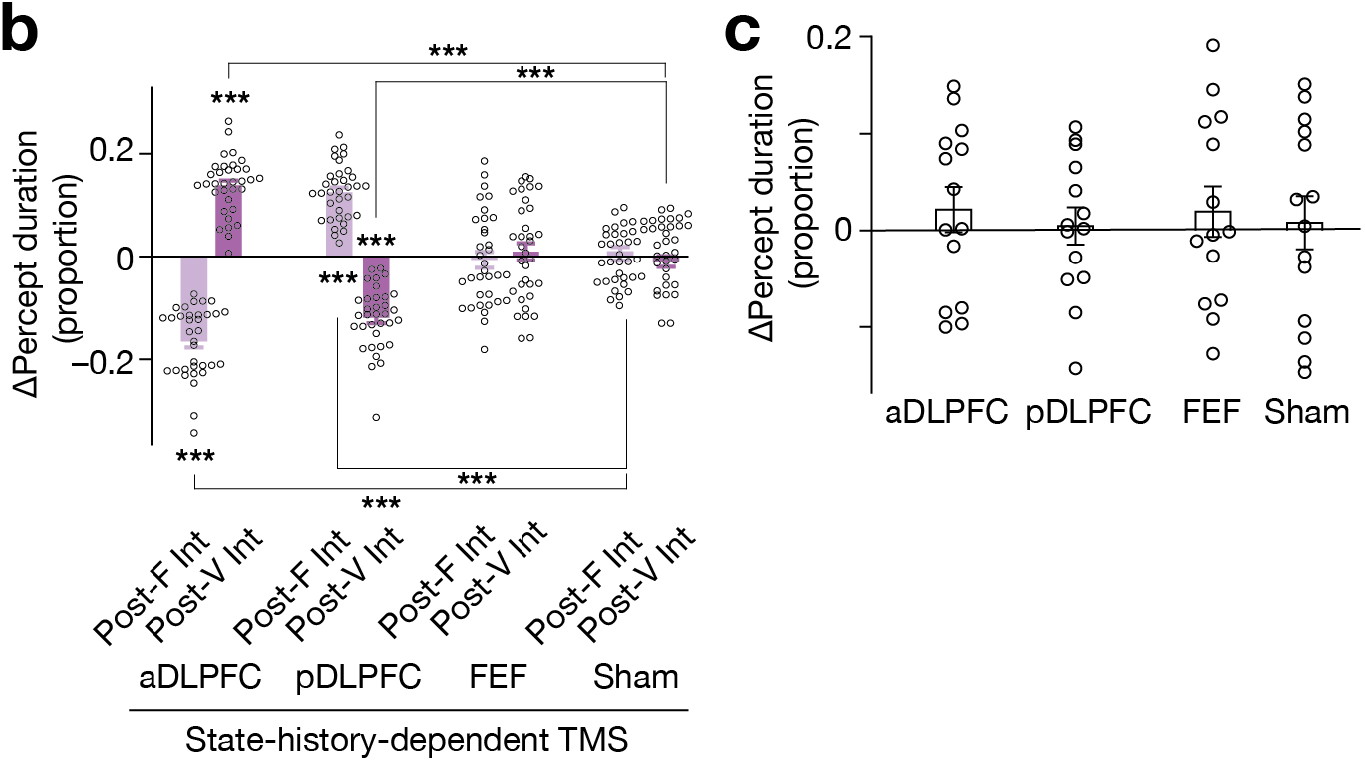
**b.** The two DLPFCs showed state-history-dependent behavioural causal effects on bistable visual perception, while FEF did not. *** indicates *P*_Bonferroni_ < 0.001 in paired *t*-tests (*df*=33). **c.** No significant behavioural change was induced after stronger and longer TMS (30-min quadripulse TMS with 90% AMT) over the same three PFC regions. The error bars show the s.e.m. The intensity of the current TMS was set at 70% AMT.

The TMS over aDLPFC during Int state immediately after F state shortened the percept duration (*t*_33_=12.9, *P*_Bonferroni_<0.001 in a post-hoc paired *t*-test, *d*=2.7), whereas aDLPFC TMS during Int state right after V state prolonged it (*t*_33_=13.3, *P*_Bonferroni_<0.001, *d*=2.3).

For pDLPFC, the Post-F Int-state-dependent TMS enhanced the perceptual stability (*t*_33_=9.1, *P*_Bonferroni_<0.001, *d*=1.6), whilst that in Post-V Int state weakened it (*t*_33_=12.7, *P*_Bonferroni_<0.001, *d*=2.3).

No change in the percept duration was observed when we administered TMS over FEF during Post-F or Post-V Int state (*t*_33_<0.58, *P*_Bonferroni_>0.05, *d*<0.19).

Given that no significant behavioural change was induced by longer and stronger TMS (here, 30-min quadripulse TMS)^34–37^ over the same PFC regions (*N*=14; *F*_2,13_=0.16, *P*=0.85 for the main effect in a two-way ANOVA; Fig. 2c), these results suggest that the causal behavioural roles of the PFC areas in the bistable visual perception become explicit and measurable only when we intervene in the neural activity in state-/state-history-dependent manners.

### Effects on energy landscape structure

These behavioural results imply that the neural mechanisms underlying this behavioural causality should be also understood in terms of brain state dynamics. Therefore, we then sought for such neurobiological accounts for these seemingly complicated behavioural observations by probing how the TMS affected the brain state dynamics.

First, we formulated working hypotheses on the neural effects by conducting numerical simulation on how TMS would affect the structure of the energy landscape that underpinned the brain state dynamics. In particular, we calculated changes in the heights of the energy barriers between the three major brain states (Fig. 2d), because the barrier heights are associated with the dwelling time in the brain states and inversely correlated with the transition frequency between them ^19^.

**Fig. 2.**
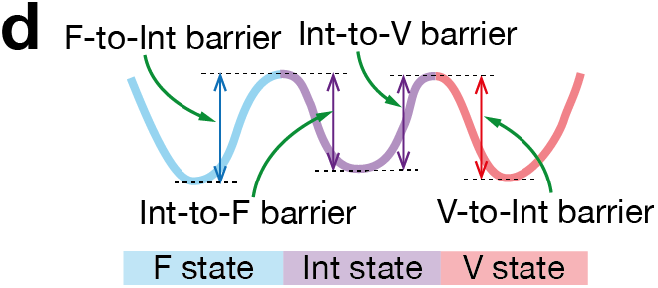
**d.** In the numerical simulation, we calculated how inhibitory TMS affects the heights of the F-to-Int, Int-to-F, Int-to-V and V-to-Int barrier for each participant.

As to aDLPFC (Fig. 2e), the numerical simulation showed that the TMS should increase the energy barrier heights between F and Int state (*F*_4,33_=340.1, *P*<10^−3^ for the main effect in a two-way ANOVA; *t*_33_>15.2, *P*_Bonferroni_<0.001 in post-hoc paired *t*-tests, *d*>2.6) and decrease the barrier height from Int to V state (*t*_33_=38.0, *P*_Bonferroni_<0.001, *d*=6.7). As a result, the ratio of the Int-to-F barrier to the Int-to-V barrier should significantly increase (*t*_33_=27.1, *P*_Bonferroni_<0.001, *d*=4.8).

**Fig. 2.**
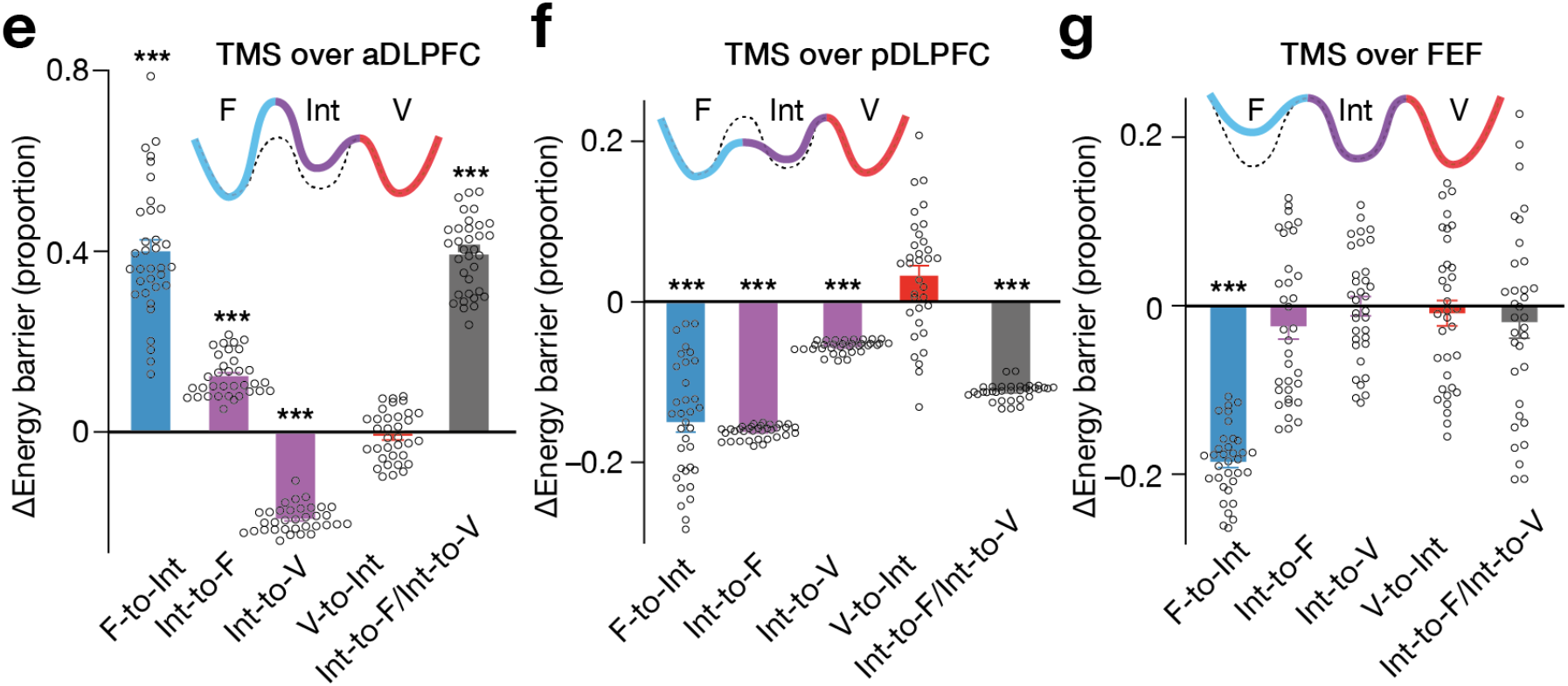
**e-g.** The numerical simulation indicated how the inhibitory TMS should affect the heights of the energy barriers between the major brain states (**panel e** for TMS over aDLPFC, **panel f** for TMS over pDLPFC and **panel g** for TMS over FEF). The solid tricolour curves schematically represent energy landscape structures after such TMS intervention, whereas the dashed curves denote the original energy landscape shown in the **panel d**. *** indicates *P*_Bonferroni_ < 0.001 in paired *t*-tests (*df*=33). The error bars show the s.e.m.

Regarding pDLPFC (Fig. 2f), the neural inhibition of the region should alleviate the energy barriers between F and Int state (*F*_4,33_=89.10, *P*<10^−3^ for the main effect in a two-way ANOVA; *t*_33_>11.6, *P*_Bonferroni_<0.001, *d*>2.0) and that from Int to V state (*t*_33_=48.9, *P*_Bonferroni_<0.001, *d*=8.6). The magnitude of the decrease in the Int-to-F barrier height should become larger than that in the Int-to-V barrier, which would result in a significant decrease in the ratio of the Int-to-F barrier to the Int-to-V barrier (*t*_33_=61.2, *P*_Bonferroni_<0.001, *d*=10.7).

For FEF (Fig. 2g), its neural suppression should lower the energy barrier from F to Int state (*F*_4,33_=32.4, *P*<10^−3^ for the main effect in a two-way ANOVA; *t*_33_=25.3, *P*_Bonferroni_<0.001, *d*=4.4) but induce no significant change in the other barriers (*t*_33_<1.7, *P*_Bonferroni_>0.05, *d*<0.27).

### Effects on brain state dynamics

These structural changes in the energy landscapes allowed us to infer the TMS-induced neural effects on the brain state dynamics (Fig. 2h).

For example, the increase in the height of the F-to-Int energy barrier—which is supposed to be induced by TMS over aDLPFC—should enhance the segregation between F and Int state, impede the transition from F to Int state and prolong the dwelling in F state. Moreover, such longer F-state dwelling should be observed in F-state-dependent TMS condition the most clearly.

**Fig. 2.**
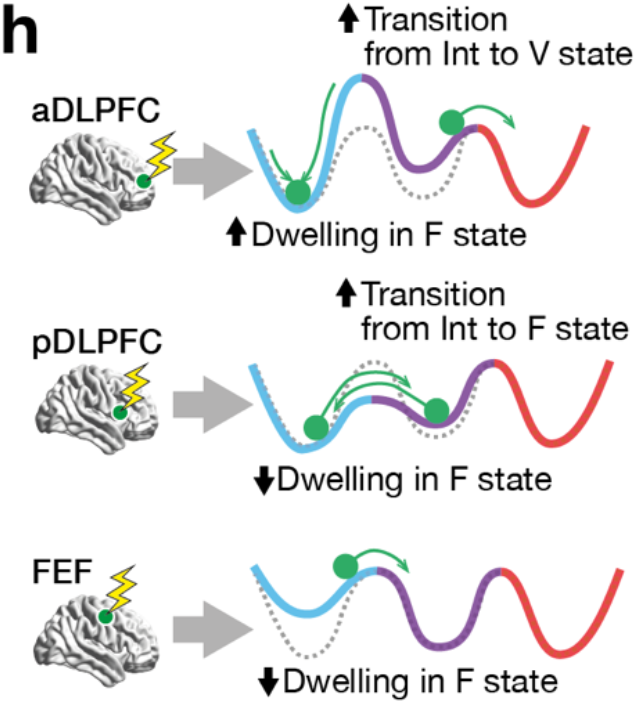
**h.** The numerical simulation (**panels e-g**) enables us to infer TMS-induced effects on the brain state dynamics. The tricolour curves schematically indicate the energy landscapes modified by TMS, whereas the dashed curves show the original energy landscapes. The green circles show the brain activity pattern: the brain state dynamics are explained as the movement of such a green ball on the energy landscape.

By the same logic, the relatively lower Int-to-V barrier, which is expected to occur in the aDLPFC TMS condition, should increase the Int-to-V transitions. Regarding the TMS over pDLPFC, the lower F-to-Int energy barrier should shorten the F-state dwelling, whereas the relatively lower barrier from Int to F state should increase the Int-to-F transitions. In the FEF TMS condition, the lower F-to-Int barrier should reduce the F-state dwelling.

We tested and confirmed these hypotheses by measuring the dwelling time of the three major brain states and transition frequencies between them in all the state-dependent TMS conditions.

In the experiments administering TMS over aDLPFC, the increase in the F-to-Int energy barrier was correlated with the longer F-state dwelling seen in the F-state-dependent aDLPFC TMS (*t*_33_=11.6, *P*_Bonferroni_<0.001, *d*=2.0; *r*_33_=0.59, *P*<0.001; Fig. 3a). Also, the relatively larger Int-to-F energy barrier was associated with more frequent Int-to-V transitions that was seen in the Int-state-dependent TMS over aDLPFC (*t*_33_=3.3, *P*_Bonferroni_<0.05, *d*=1.8; *r*_33_=0.52, *P*<0.001; Fig. 3b).

**Fig. 3.**
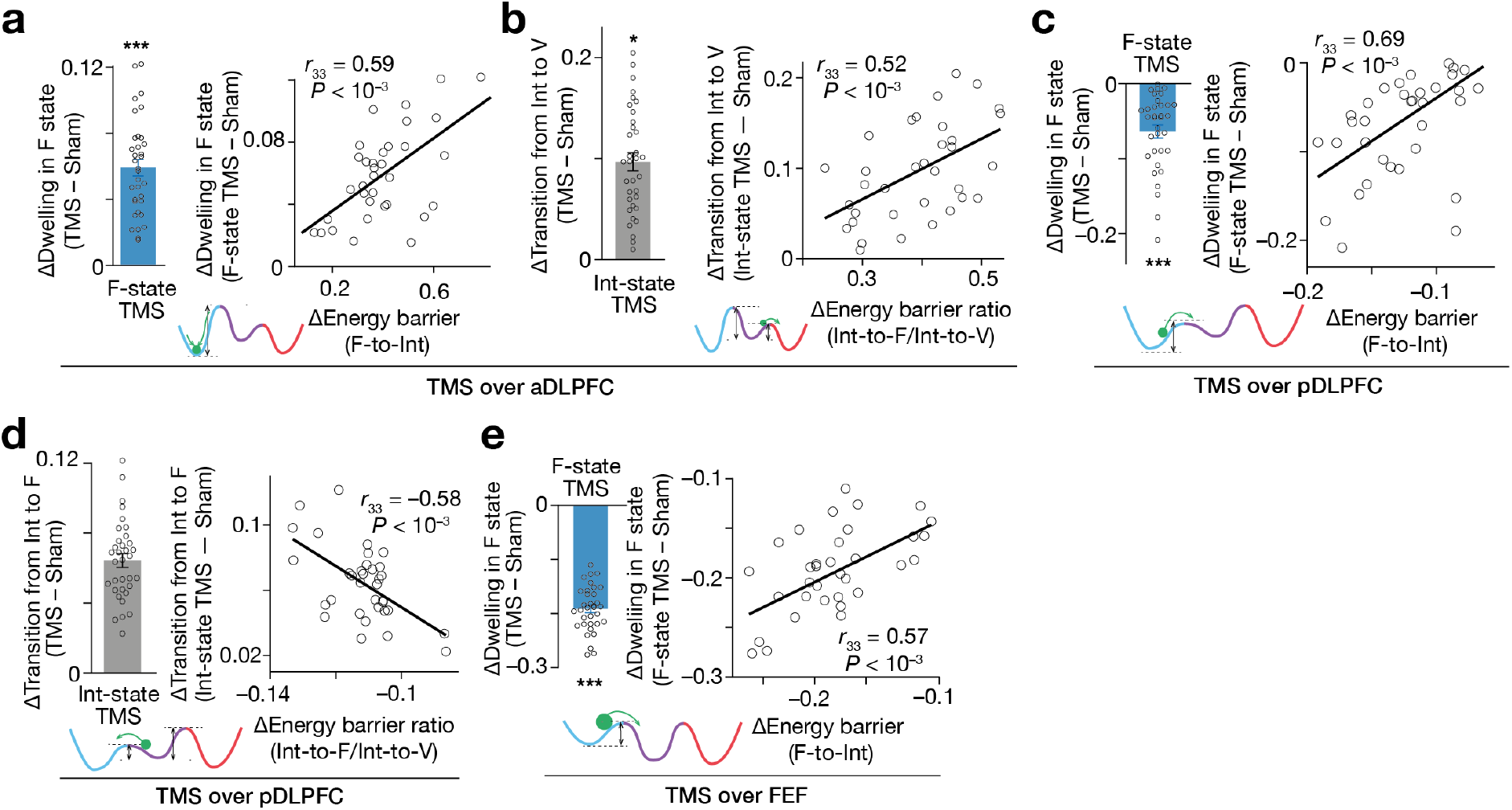
**a-e.** The hypotheses presented in **Fig. 2h** were confirmed. The X-axes show the changes in the hypothetical energy barrier heights, whereas the Y axes indicate TMS-induced neural effects on the brain state dynamics. The circles represent the participants. *** and * indicate *P*_Bonferroni_ < 0.001 and *P*_Bonferroni_ < 0.05 in paired *t*-tests (*df*=33), respectively. The error bars show the s.e.m.

In the sessions administering TMS over pDLPFC, the lower F-to-I energy barrier was correlated with the shorter F-state dwelling that was measured in the F-state-dependent neural suppression (*t*_33_=6.9, *P*_Bonferroni_<0.001, *d*=1.2; *r*_33_=0.69, *P*<0.001; Fig. 3c). The relatively lower Int-to-F barrier accurately predicted more frequent Int-to-F transitions (*t*_33_=13.0, *P*_Bonferroni_<0.001, *d*=2.8; *r*_33_=–0.58, *P*<0.001; Fig. 3d) that were seen in the Int-state-dependent TMS condition.

In the FEF conditions, the lower F-to-Int barrier was associated with the shorter F-state dwelling that was observed in the F-state-dependent TMS session (*t*_33_=26.1, *P*_Bonferroni_<0.001, *d*=4.8; *r*_33_=0.57, *P*<0.001; Fig. 3e).

These results clarify TMS-induced effects on the brain state dynamics during the bistable visual perception and demonstrate that such neural effects are underpinned by the structural changes in the hypothetical energy landscapes.

### Temporal decay of neural effects

How long did such neural effects continue after each TMS? Instinctively, they should not last significantly longer compared to the intervals between the neural stimulations (~10sec). If the current state-dependent TMS could induce as long neural effects as conventional repetitive TMS protocols, it would be difficult to find sound reasons why the 30-min continual TMS did not affect the perceptual stability behaviourally (Fig. 2c) but the state-dependent TMS did (Figs. 2a and 2b).

We investigated this question by tracking the magnitudes of the correlations between the neural effects and the energy barrier changes (Figs. 3a-e) when we were sliding the time window that was used to quantify the neural effects (the upper panel of Fig. 3f). This analysis detected that all the correlations began to weaken after an approximately 1.5-sec mild decline (the lower graph in Fig. 3f). This result indicates that the current TMS-induced neural effects started to decay within ~1.5sec after the stimulation; thus, the brain state dynamics and corresponding energy landscapes began to return to the original forms in such a time scale.

**Fig. 3.**
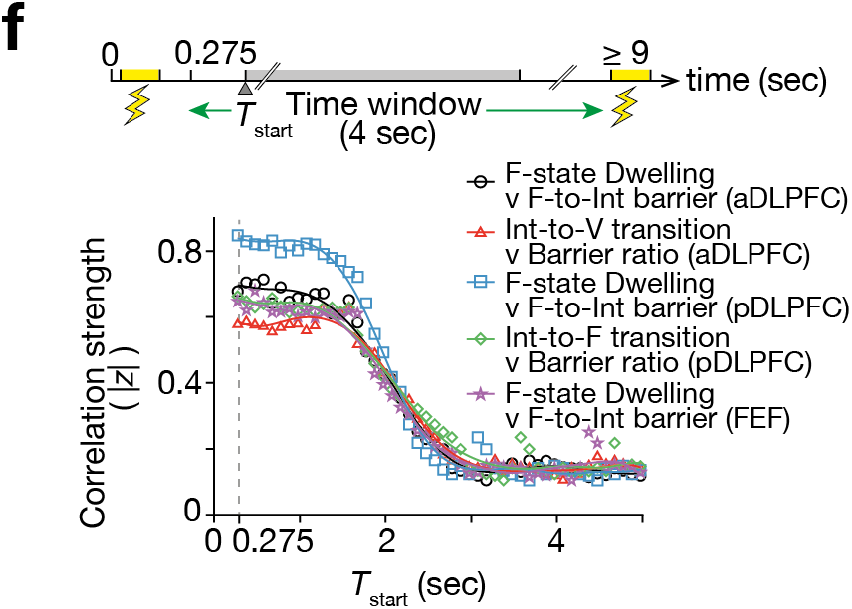
**f.** To assess the duration of the TMS-induced neural effects, we examined how the correlations seen in **panels i-m** changed when we moved the 4-sec time window that was used to measure the neural effects. In all the correlations, the correlation strength (here, the absolute value of the Fisher-transformed correlation coefficient, |*z*|) began to decay ~1.5-sec after the TMS. The X-axis shows the start timing of the 4-sec time window (*T*_start_).

### Neural mechanisms behind behavioural causality

Finally, by evaluating relationships between these behavioural, numerical and neural responses with mediation analysis, we clarified that the causal behavioural effects are induced by transient changes in the brain state dynamics and attributable to structural changes in the energy landscapes (*P*<0.05 for all the indirect effects, *α*×*β*; Figs. 4a-4g).

**Fig. 4.**
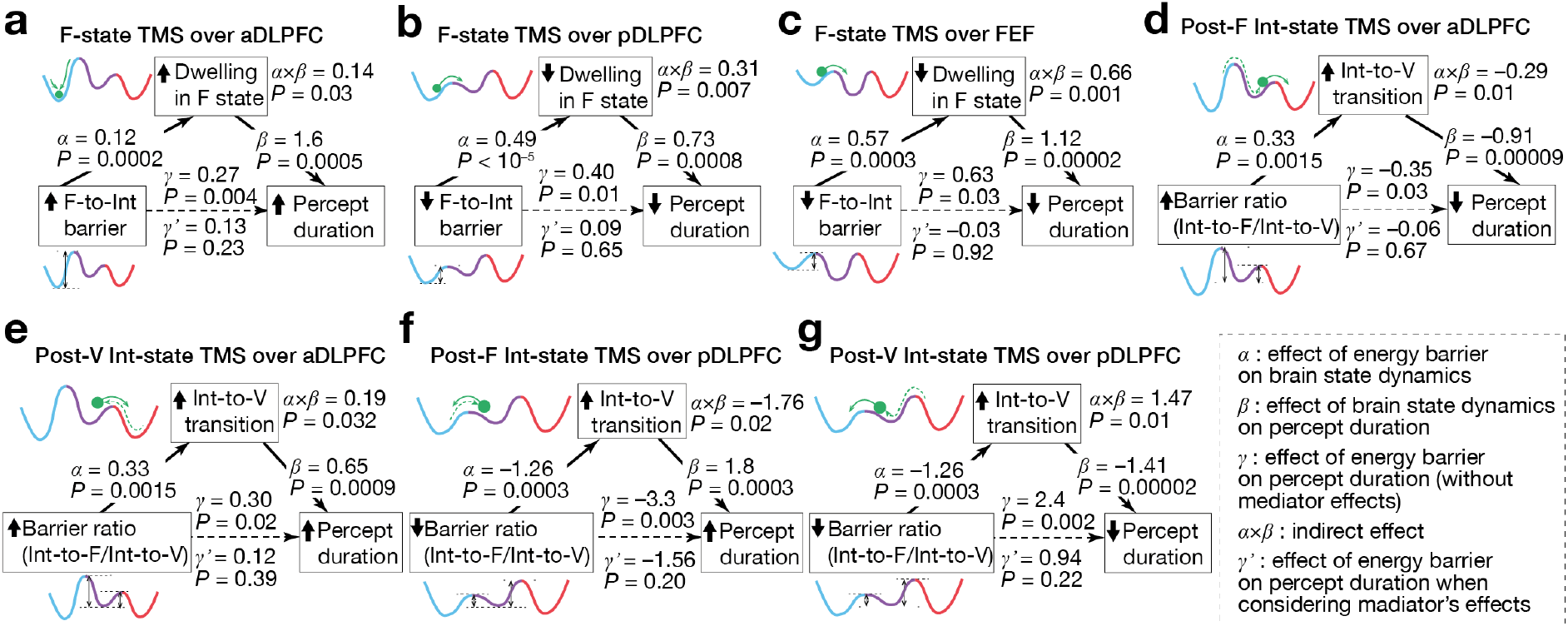
**a-g.** The mediation analyses showed that the structural changes in the energy landscapes affect the brain state dynamics, which results in the causal behavioural responses. The statistical significance of the *α*, *β* and *γ* validate our application of the mediation analysis to the current data. The statistical significance of the *α*×*β* and the insignificance of the *γ*’ support the conclusions.

As to the state-dependent behavioural changes, the longer percept duration seen after the F-state-dependent TMS over aDLPFC was largely due to the prolonged F-state dwelling, which was induced by the higher energy barrier from F state to Int state (Fig. 4a). In contrast, the de-stabilised visual perception induced by the F-state-dependent TMS over pDLPFC/FEF was attributable to the shorter F-state dwelling time, which was caused by the lower F-to-Int energy barrier (Figs. 4b and 4c).

If we admit that the percept duration is closely linked with the length of the F-Int-V travel (Fig. 1h), the state-history-dependent behavioural causality could also be seen as a consequence of the transient changes in the brain state dynamics.

In the mediation analysis, the shorter percept duration observed after the Post-F Int-state-dependent TMS over aDLPFC was caused by the increase in the Int-to-V transitions, which was induced by the larger gap between the Int-to-F barrier and Int-to-V barrier (Fig. 4d). This statistical result is reasonable because the relatively lower Int-to-V energy barrier would facilitate the Int-to-V transitions transiently, accelerate the completion of the F-Int-V travel and shorten the percept duration.

Conversely, the longer percept duration yielded by the Post-V Int-state-dependent TMS over aDLPFC is interpretable as a result of the relatively higher Int-to-F energy barrier’s impeding the Int-to-F transitions, accelerating the backward moves to V state and slowing down the completion of the F-Int-V travel (Fig. 4e).

The mediation analysis showed that the state-history-dependent causal roles of pDLFPC were also explainable by the same logic. The longer percept duration yielded by the pDLPFC TMS during Post-F Int state could be regarded as a behavioural manifestation of the temporary slowdown of the F-Int-V travel due to the more frequent backward Int-to-F transitions, which was originated from the relatively lower Int-to-F energy barrier (Fig. 4f). In contrast, the shorter percept duration induced by the pDLPFC TMS during Post-V Int state can be seen as results of the acceleration of the F-Int-V travel due to the more frequent forward moves from Int to F state, which was induced by the lower Int-to-F energy barrier (Fig. 4g).

## Discussion

This study has identified state-/state-history-dependent causal behavioural roles of the three PFC regions—right FEF and anterior/posterior DLPFCs—during the SFM-induced bistable visual perception. The behavioural causality was determined by the large-scale brain state dynamics which was underpinned by the hypothetical energy landscape structures. Moreover, the current findings have clarified distinct functions of the PFC regions in the brain state dynamics: the activation of aDLPFC enhances the mutual interaction and functional integration between Frontal-area-dominant and Intermediate state, whereas pDLPFC activity supports the functional diversity between the two PFC-active brain states; the FEF activation plays a critical role in the stabilisation of Frontal-area-dominant state.

These findings may not be directly applicable to other types of multistable visual perception, such as binocular rivalry, which is often explained by neural responses in lower-level brain architectures including the visual cortex ^31,32^ and lateral geniculate nucleus ^33^. Instead, for the SFM-induced bistable perception, the current behavioural observations were robust. All these behavioural findings were replicated in a small but independent cohort (*N*=14; *t*_13_>2.9, *P*<0.01 in one-sample *t*-tests; Fig. 5a). Moreover, another independent experiment (*N*=15) showed that, as in our previous work ^35,36^, the excitatory neural modulation induced behavioural effects opposite to those yielded by the inhibitory one (*t*_14_>2.8, *P*<0.01 in one-sample *t*-tests; Fig. 5b), which added empirical support for the current observations.

**Fig. 5.**
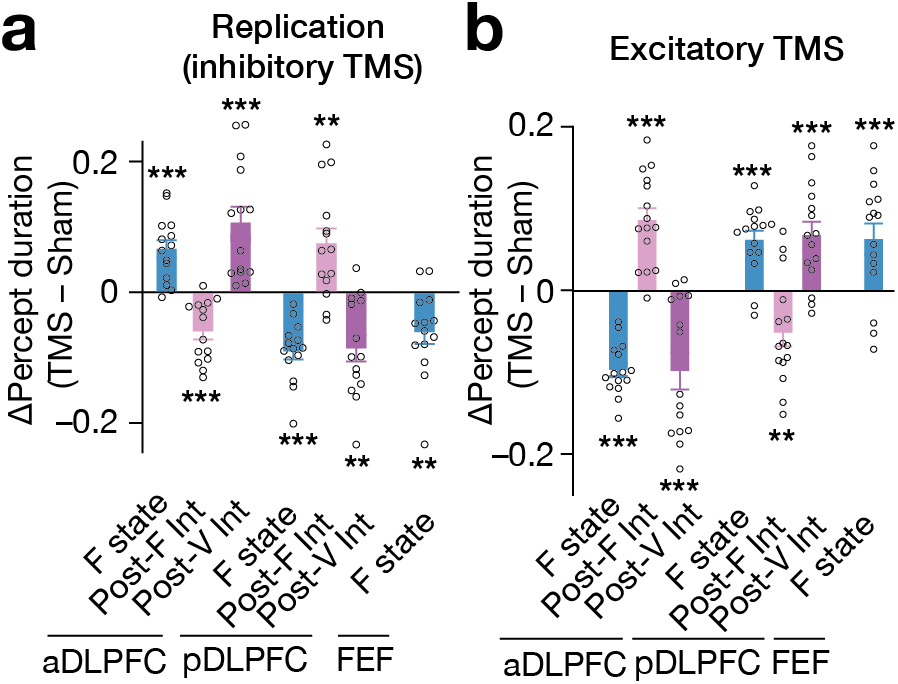
**a.** All the TMS-induced significant behavioural changes seen in the main experiment were replicated in an independent cohort (*N*=14). **b.** We conducted another validation experiment by administering excitatory TMS over the same PFC regions of another independent cohort (*N*=15). The excitatory stimulated induced behavioural effects opposite to those seen in the inhibitory TMS experiments. Each circle represents each participant. *** and ** indicate *P*<0.001 and *P*<0.01 in one-sample *t*-tests (*df*=13 for **Fig. 5a** and *df*=14 for **Fig. 5b**), respectively. The error bars show the s.e.m.

The current observations re-highlight the fact that brain-behaviour causality is changing so dynamically even during a simple cognitive task that we often cannot behaviourally detect it in conventional neurostimulation experiments that do not consider the temporal changes of the brain states ^21,38^. Previous work has addressed this issue by controlling, inferring or monitoring the brain state: some studies controlled the brain state using external stimuli and applied the TMS when a specific sensory stimulus was presented to the participants ^39–41^; a clinical research adopted emotional states as indicators of the neural activity and determined the timing of the deep brain stimulation based on such inference ^42^; neurophysiological studies monitored neural activity and determined the timing of brain stimulation based on the frequency, phase or power of the neural signal ^28–30,43,44^.

The current TMS method is categorised into the last group and could be seen as advancement of such direct-neural-monitoring-based brain stimulation. Differently from the previous work monitoring a single neural activity ^28–30,43,44^, the TMS system used here can track the brain state using neural activity patterns recorded from multiple remote brain regions. Given the multi-regional brain state dynamics underpin complex cognitive activities ^19,23,45^, such a multivariate monitoring approach could be regarded as a more effective manner to investigate more physiological brain-behaviour causality.

Moreover, the combination of EEG-triggered TMS and energy landscape analysis may become foundation of a novel tool to control seemingly unstable behaviours. In fact, two-month longitudinal experiments (*N*=63) revealed accumulative effects of this closed-loop neural modulation system. Despite its weak power—approximately 78% of the power of the similar TMS protocol ^35,36^—, the behavioural effects became larger along with the weekly TMS sessions (*t*_12_>3.2, *P*_Bonferroni_<0.05 in paired *t*-tests; Fig. 6a). In addition, the F-state-dependent TMS affected the baseline perceptual stability, which were observed even one-week after the two-month TMS experiments (*F*_6,12_=16.4, *P*<10^−3^ for the main effect in a two-way ANOVA; *P*_Bonferroni_<0.01 in post-hoc one-sample *t*-tests; Fig. 6b). Considering that the perceptual stability seen in this bistable perception test was relevant to autistic cognitive rigidity ^46^, these longitudinal observations may become a basis for new clinical non-invasive neural interventions and accelerate the development of such state-dependent brain stimulation methods.

**Fig. 6.**
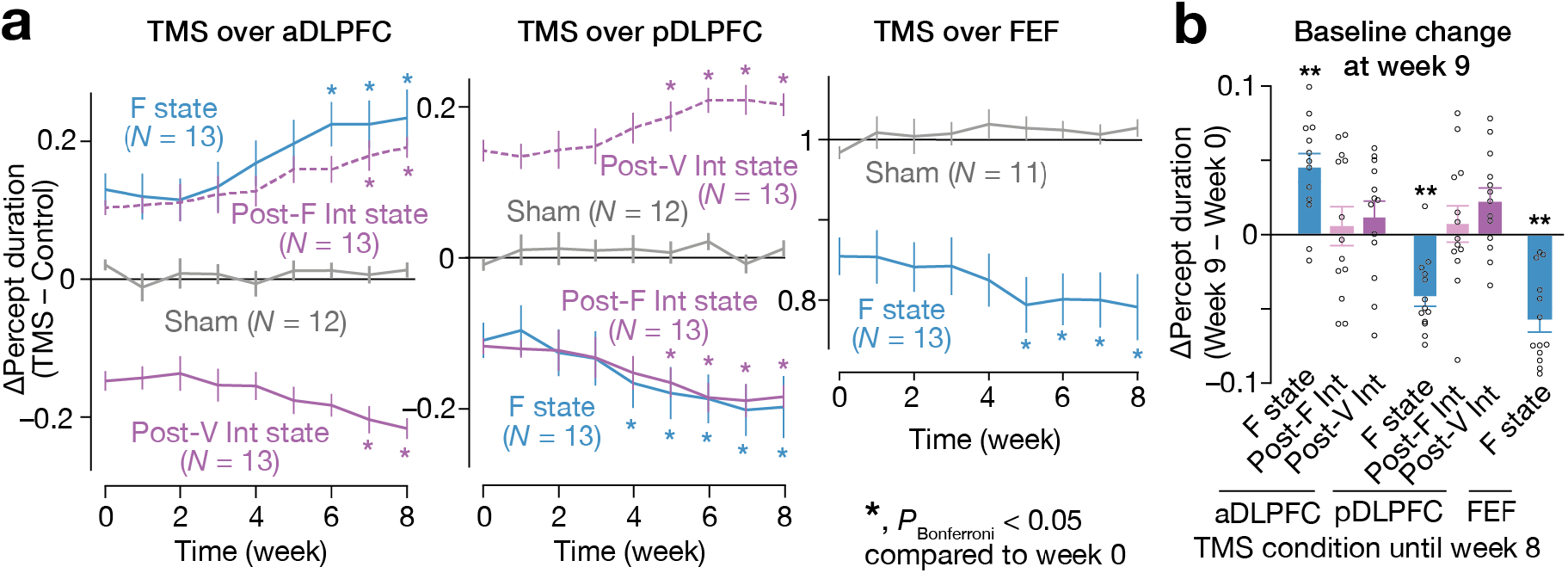
**a-b.** We examined the accumulative effects of the state-/state-history-dependent TMS on the bistable visual perception in two-month longitudinal experiments. We focused on the seven TMS conditions that induced significant behavioural effects (**Fig. 2a and 2b**). In parallel, we also examined behavioural changes in three sham conditions in which TMS was applied over either of the three PFC regions with random intervals. In all the TMS conditions, the participants became more responsive to the TMS (**panel a**). Even after the two-month weekly TMS experiments, we observed significant changes in the baseline percept duration in three of the seven TMS conditions (**panel b**). In **panel a**, the Y-axes show the changes in the median percept duration (TMS – Control), and * indicates *P*_Bonferroni_ < 0.05 in paired *t*-tests with comparisons to the week-0 values (*df*=12). In **panel b**, the Y-axis indicates the changes in the control performance between week 0 and 9. ** indicates *P*_Bonferroni_ < 0.01 in one-sample *t*-tests (*df*=12).

To interpret the current observations in psychological contexts such as predictive coding ^2,11^, more model-based neuroimaging studies and theoretical work would be necessary. However, this study has resolved the long-lasting controversy over prefrontal causal roles in multistable perception ^2^ and revealed distinct functions of the PFC in the brain state dynamics that underpin spontaneous perceptual inference. Furthermore, the combination of the brain-state tracking method and neural-activity-dependent brain stimulation system may re-ignite neurobiological investigation on state-dependent dynamic causality ^20^ in human cognition and become another foundation of more effective neural perturbation tools to intervene in neuropsychiatric conditions.

## Materials and Methods

### 1.1. Overall study design

This study consisted of seven experiments and one numerical simulation (Supplementary Fig. 1).

First, 65 healthy adult individuals underwent two control experiments, in which we recorded their EEG signals while they were experiencing bistable visual perception induced by a structure-from-motion (SFM) stimulus (Fig. 1a). In the Control Experiment I, The EEG data were used to (i) identify individual brain dynamics during bistable visual perception and (ii) verify the locations of EEG electrodes and TMS stimulation site. In the Control Experiment II, we examined the accuracy of the brain-state tracking and state-dependent TMS system.

Second, 35 of all the 65 individuals participated in the main experiment. In this experiment, using the information about individual brain state dynamics, we administered inhibitory TMS to the participants in state-/state-history-dependent manners. In parallel, we conducted a numerical simulation to predict how inhibitory TMS changed individual energy landscape structures and, resultantly, brain state dynamics. Thirty four of the 35 participants completed this experiment. This sample size of the main experiment was determined on a power analysis (effect size = 0.5; power = 0.8; alpha = 0.05 for paired *t*-tests).

Other 15 participants underwent a replication experiment that re-tested behavioural causality found in the main experiment, and 14 of them completed the experiment. Afterwards, the 14 participants underwent a conventional 30-min quadripulse TMS (QPS) ^34–36^ experiment, in which they received the QPS over one of the three PFC regions (aDLPFC, pDLFPC and FEF) during rest.

The other 15 individuals participated in a validation experiment that examined whether excitatory TMS induced behavioural effects opposite to those caused by inhibitory TMS.

The 63 individuals who completed either of these experiments participated in a longitudinal experiment, which evaluated the accumulative effects of the current neural stimulation method.

**Supplementary Fig. 1.**
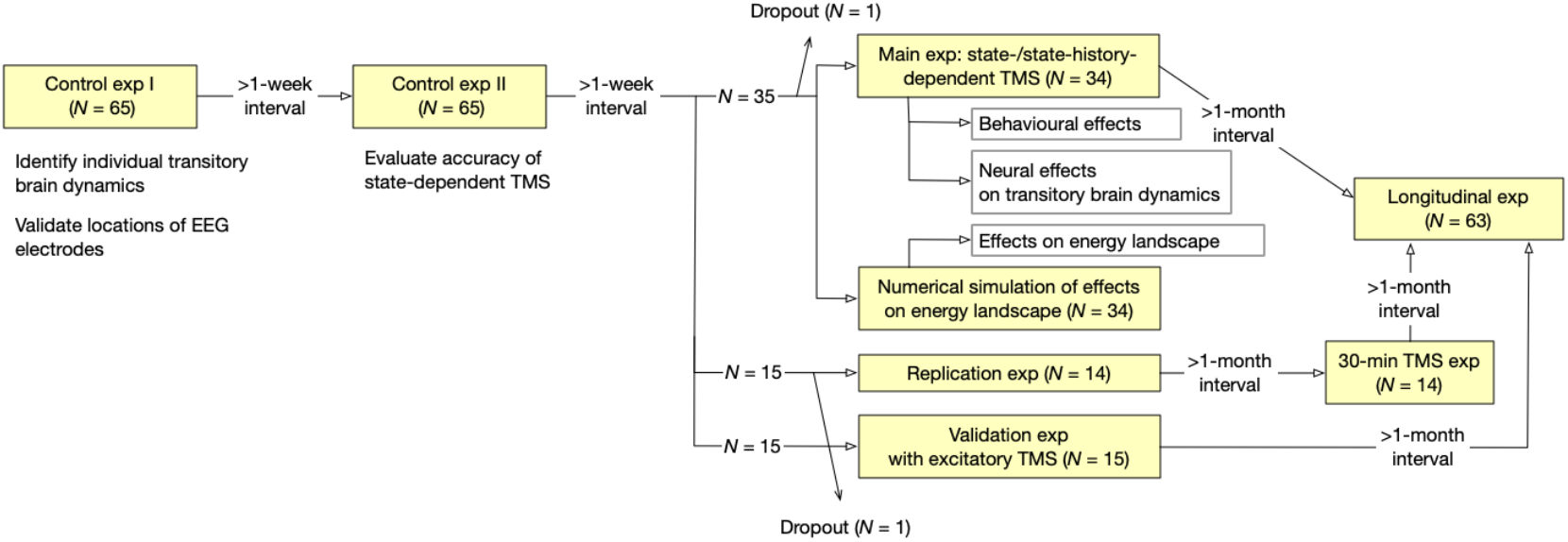
The current study consisted of seven EEG/TMS experiments and one numerical simulation.

### 1.2. Participants and ethics

All the 65 participants were right-handed adults (Edinburgh Handedness Inventory laterality score = 75±16, mean±sd). None of them had neurological, psychiatric or other medical history and was free from any contraindication to TMS experiments ^47^.

Except for the control experiments and 30-min QPS experiment, all the experiments asked the participants to undergo the tests continually over several weeks; therefore, some of the participants did not complete the study allegedly due to their busyness of life. Consequently, the final sample size was 34 for the main experiment (age = 23.1±1.8; female = 12), 14 for the replication experiment (age = 22.2±2.0; female = 8), 15 for the validation experiment using excitatory TMS (age = 22.7±1.7; female = 7) and 63 for the longitudinal study (age = 22.8±1.9; female = 27). Including the participants who dropped out, no participant reported adverse effects throughout this study.

This study was approved by Institutional Ethics Committees in RIKEN and The University of Tokyo. The TMS protocols used here complied with the guideline issued by the Japanese Society for Clinical Neurophysiology and that by International Federation of Clinical Neurophysiology ^48^. All the participants provided written informed consents before any experiment and were financially compensated for their participation.

### 2.1. Device setup: test of bistable visual perception

The test design of bistable visual perception paradigm in this study is essentially the same as that used in our previous work ^19,46^. The participants were presented with a structure-from-motion (SFM) stimulus (Fig. 1a), a sphere consisting of 200 sinusoidally moving white dots in a black background (angular velocity, 120°/sec) with a fixation cross (0.1° × 0.1°) at the centre of the 27-inch LCD monitor (BenQ PD2710, resolution: 2560×1440).

In each run, the participants were instructed to see the SFM stimulus for 90 sec with their chins put on a chin rest. They were asked to push one of the three buttons according to their visual perception: one for upward rotation, another for downward rotation, and the other for unsure or mixture perception. After sufficient training sessions, the participants repeated this run six times in all the experiments except for the control one. The stimulus presentation and response recording were conducted with PsychToolbox 3 in MATLAB (MathWorks, Inc).

The proportion of the mixture perception was sufficiently small in all the participants, all the experiments (1.4±0.5% of all stimulus presentation times, mean±sd). Thus, we focused on the time during which participants were clearly aware of the direction of the rotation. For each participant, we measured the duration of the clear perception and calculated the median of the duration to evaluate their perceptual stability. The median duration was adopted because the perceptual durations showed long-tailed distributions.

### 2.2. Device setup: EEG

Throughout entire this study, we recorded EEG signals from seven regions of interest (ROIs) to monitor brain state transitions using a TruScan Research EEG system with 32 TMS compatible Ag/AgCl ring electrodes (Deymed Diagnostic, Czech Republic).

The seven ROIs consisted of the right FEF (x = 38, y = 0, z = 60 in MNI coordinates), aDLPFC (x = 44, y = 50, z = 10), pDLPFC (x = 48, y = 24, z = 9), anterior superior parietal lobule (aSPL; x = 36, y = –45, z = 44), posterior superior parietal lobule (pSPL; x = 38, y = −64, z = 32), lateral occipital complex (LOC; x = 46, y = −78, z = 2) and V5 (hMT/V5; x = 47, y = −72, z = 1) (Fig. 1b). All the ROIs were selected from a line of previous studies on bistable visual perception ^4,5,16,49–52^. To track the same neural transitions in the same brain dynamics as those seen in our previous work ^19^, we adopted the same MNI coordinates as those used in the study.

Using a stereoscopic neuro-navigation system (Brainsight Neuronavigation, Rogue Research, UK) and structural MRI brain images, we located the TMS-compatible EEG electrodes right above on these seven ROIs. Also, for the following calculation of Hjorth signals ^53^(see Section 3.2.1), we put three other EEG electrodes around each ROI electrode (i.e., four electrodes were used for one ROI; Supplementary Fig. 2). The other four electrodes (i.e., 32 electrodes – 4 electrodes/ROI × 7 ROIs = 4 electrodes) were located on Fpz, Oz, A1 and A2 in accordance with 10-20 international system (Seeck et al. 2017). These electrodes were firmly placed on the heads of the participants with elastic caps

After confirming that the impedance was less than 5kΩ in all the electrodes, we recorded the EEG signals with a 32-channel amplifier (TruScan EEG LT 32ch Headbox, Deymed Diagnostic; 6kHz of analogue sampling frequency), in which the signals underwent low-pass filtering (cut-off frequency: 1.25kHz) and were down-sampled to 3kHz at almost simultaneously (latency <5ms).

**Supplementary Fig. 2.**
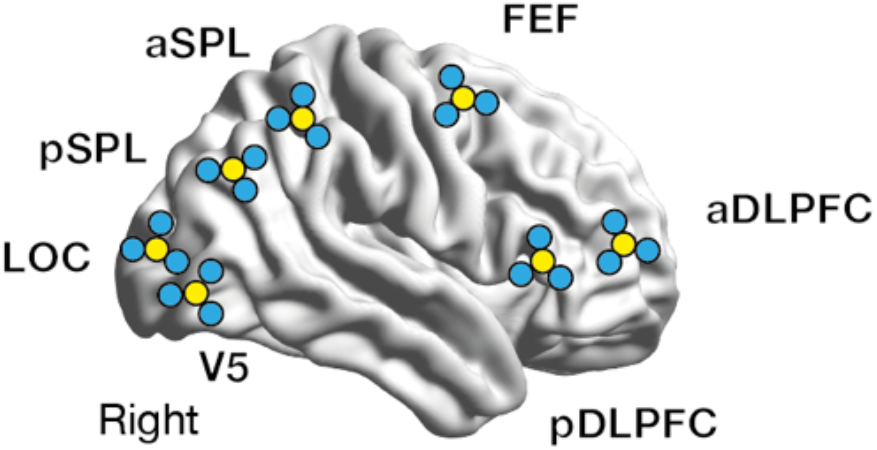
We placed TMS-compatible EEG electrodes in an original way. First, seven electrodes were located right above on the seven ROIs using a stereoscopic neuro-navigation system. Then, for each ROI, three additional electrodes were placed right around the electrodes for the following Hjorth signal calculation. The yellow circles schematically represent the electrodes put above the ROIs, whereas the blue ones indicate the neighbouring electrodes.

### 2.3. Device setup: TMS

To deliver both inhibitory and excitatory stimulation, we used a quadripulse TMS system that can deliver a train of four monophasic magnetic pulses at once from interconnected four magnetic stimulators (70mm-diameter double coil, DuoMag MP, DeyMed Diagnostic, Czech Republic).

To reduce the length of the after-effects of the TMS and focus on transient causality, our EEG-triggered TMS system did not use conventional quadripulse TMS protocols ^34,35,37,54,55^ in which the TMS was continually administered throughout 30min and whose after-effects are considered to last for more than 60min. Instead, in the main experiment, we conducted a single set of four-pulse 20Hz monophasic TMS for neural suppression, whereas a set of four-pulse 200Hz monophasic TMS was administered as an excitatory stimulation in the validation study.

In addition, to obtain sufficient length of clean EEG data for the following brain-state tracking, we put ≥9-sec intervals between the stimulations and set the intensity of the TMS stimulation at a lower level (70% of active motor threshold, AMT) compared to that in the conventional quadripulse TMS protocol (90% of AMT)^35,36^. In EEG-triggered TMS, such weak stimulations were considered to induce sufficiently large neural effects ^28^.

The AMT was calculated based on motor evoked potentials (MEPs) of the right first dorsal interosseous (FDI) muscle of each participant in prior single-pulse TMS experiments: the AMT was set as the lowest TMS intensity that evoked a small response (>100μV) when the participants maintained a slight contraction of the right FDI (approximately 10% of the maximum voluntary contraction) in more than the half of 10 consecutive trials ^35,36^.

The MEPs of the right FDI were recorded using a pair of 9mm-diameter Ag/AgCl surface cup electrodes that were placed over the muscle belly and the metacarpophalangeal joint of the index finger ^56^. Before offline analyses, the MEP signals underwent a temporal filter (100Hz–3kHz). The optimal place for the single-pulse TMS for the right FDI muscle was determined as the area over which the simulation induced the largest MEP.

The mean AMT across the entire participant cohort in the three types of the current TMS experiment was 35.8±7.9% of the maximum stimulator output.

### 3.1. EEG Analysis setup

In this study, we analysed the EEG data in both offline and online manners. We described the details of the two types of EEG analysis in the following sections.

#### 3.2.1. Offline EEG analysis: preprocessing

The offline EEG analysis was applied to data obtained in the control experiment. We conducted the following conventional preprocessing ^57^ to the EEG data using MATLAB (MathWorks, US) and EEGLAB ^58^.

First, the EEG data were referenced to the average across all the electrodes, down-sampled to 300Hz and underwent a temporal filter (1–80Hz). Then, we conducted an independent component analysis to remove cardio-ballistic artefacts and other artefacts induced by eye blinks, eye movements and muscle activity. Next, we marked epochs whose mean global field power was too large (> 5SD of mean power across entire recording) and excluded those time periods in all the following main analysis. We then filtered the data to delta (1–4Hz), theta (4-8Hz), alpha (8–13Hz), beta (13–30Hz) and gamma (30–80Hz) bands and estimated a Hilbert envelope amplitude for the gamma-band signal^57,59^.

We used the Hilbert envelope amplitude for the gamma band as a neural signal for each electrode, because the aim of the current EEG recording was to trace the brain state dynamics that was seen in our previous fMRI study ^19^ and the gamma-band signal dynamics were correlated with fMRI signals ^57,59^. Finally, after removing autocorrelation, we calculated a Hjorth signal for each ROI in the following sum-of-difference manner ^53^: Hjorth signal for ROI_*i*_ = (electrode just above ROI_*i*_ – surrounding electrode 1) + (electrode just above ROI_*i*_ – surrounding electrode 2) + (electrode just above ROI_*i*_ – surrounding electrode 1).

#### 3.2.2. Offline EEG analysis: fitting of pairwise maximum entropy model

We then conducted the energy landscape analysis ^19,23–26^ of the preprocessed datasets of the seven ROIs. For each participant, the EEG signals were concatenated across different runs.

First, as in our previous fMRI work on the brain dynamics during bistable perception, we binarised each ROI time-series data using the temporal average of the signals as the thresholds. We then fitted a pairwise maximum entropy model (MEM) to the seven. binary time-series signals in the same manner as in our previous studies ^19,23^.

A signal pattern of the seven ROIs at time *t* was described such as 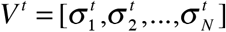 where 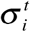 represents a binary activity of ROI_*i*_ at time *t* (i.e., 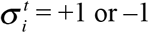) and *N* is the number of the ROIs (i.e., here, *N* = 7). In the principle of maximum entropy ^60^, the average ROI activity 〈σ_*i*_〉 and average pairwise interaction 〈σ_*i*_σ_*j*_〉 are constrained by the empirical data, and the appearance probability *P*(*V_k_*) of an ROI activity pattern *V_k_* should obey Boltzmann distribution, because such a distribution maximises the information entropy. Therefore, the appearance probability of the activity pattern *V_k_* can be stated as 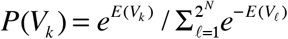, where 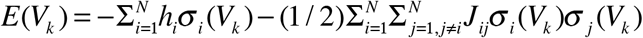. Here, σ_*i*_(*V_k_*) is the binary activity of ROI_*i*_ in the activity pattern *V_k_*, whereas *h_i_* represents the basal activity of ROI_*i*_ and *J_ij_* indicates a pairwise interaction between ROI_*i*_ and ROI_*j*_. Using the *P*(*V_k_*), we calculated the model-based mean ROI activity 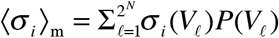 and model-based mean pairwise interaction 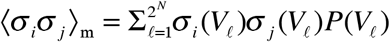. Using a gradient ascent algorithm, we adjusted *h_i_* and *J_ij_* until these 〈σ_*i*_〉_m_ and 〈σ_*i*_〉_m_ were approximately equal to the empirically obtained 〈σ_*i*_〉 and 〈σ_*i*_σ_*j*_〉.

The accuracy of this MEM fitting was examined by estimating a Pearson correlation coefficient between the model-based appearance probability and empirically-obtained appearance probability and calculating a proportion of Kullback-Leibler divergence in this 2^nd^-order model (*D*_2_) to that in the 1^st^-order model (*D*_1_) as follows ^19,23,61^: (*D*_1_ − *D*_2_)/*D*_1_. In the control experiment, the Pearson correlation was larger than 0.95 and the KL-divergence-based accuracy was larger than 84%.

#### 3.2.3. Offline EEG analysis: disconnectivity graph in energy landscape analysis

Next, we built an energy landscape and searched for major brain states. The energy landscape was defined as a network of brain activity patterns *V_k_* (*k* = 1, 2, …, 2^*N*^) with their energy *E*(*V_k_*), in which two activity patterns were regarded as adjacent if and only if they took the opposite binary activity at a single ROI. We then searched for local energy minima, whose energy values were smaller than those of all the *N* adjacent patterns.

We then examined hierarchal structures between the local minima by building disconnectivity graphs as follows ^19,23^: (i) first, we prepared a so-called hypercube graph, in which each node representing a brain activity pattern was adjacent to the *N* neighbouring nodes. (ii) Next, we set a threshold energy level, *E*_threshold_, at the largest energy value among the 2^*N*^ nodes. (iii) We then removed the nodes whose energy values were ≥*E*_threshold_. (iv) We examined whether each pair of local minima was connected by a path in the reduced network. (v) We repeated steps (iii) and (iv) after moving *E*_threshold_ down to the next largest energy value. We ended up with a reduced network in which each local min was isolated. (vi) Based on the obtained results, we built a hierarchical tree whose leaves (i.e., terminal nodes down in the tree) represented the local minima and internal nodes indicated the branching points of different local minima.

#### 3.2.4. Offline EEG analysis: structure of energy landscape

Based on this dysconnectivity graph, we then estimated basin sizes of the local minima as follows. We first chose a node *i* from the 2^*N*^ nodes. If any of its neighbour nodes had a smaller energy value than the node *i*, we moved to the neighbour node with the smallest energy value. Otherwise, we did not move, indicating that the node was a local min. We repeated this protocol until we reached a local min. The initial node *i* was then assigned to the basin of the local min that was finally reached. This classification procedure was repeated for all the 2^*N*^ nodes. The basin size was defined as the fraction of the number of the nodes belonging to the basin.

The energy barrier between local minima *ℓ* and *m* was defined based on the procedure of building the disconnectivity graph. When we built the disconnectivity graph by lowering the threshold energy level *E*_threshold_, we searched for the lowest *E*_threshold_ at which the two local minima were still connected. The height of the energy barrier from local min *ℓ* to *m* was then defined as the difference between the *E*_threshold_ and the energy value for the local min *ℓ*, whereas that from local min *m* to *ℓ* was defined as the difference between the *E*_threshold_ and the energy value for the local min *m*.

Then, as in our previous work ^19^, we summarised the local minima as follows: if the energy barriers from the local min *ℓ* to *m* was lower than a threshold (=1, here) and the energy value of the local min *ℓ* was larger than that of local min *m*, we regarded the local min *ℓ* and its basin as elements of the basin of the local min *m*. By repeating this procedure, we found that in all the participants their brain activity patterns during bistable visual perception could be classified into any of the three major basins, which corresponded to Frontal-area-dominant, Visual-area-dominant and Intermediate states. The energy barrier threshold for this summarisation was set at the same value as in our previous work ^19^.

Through this coarse-graining procedure, we defined the three major brain states (i.e., Frontal, Intermediate and Visual states) and calculated structural indices of the energy landscapes (i.e., the height of the energy barrier). In addition, this summarisation allowed us to classify all the nodes (i.e., brain activity patterns) on the energy landscape—except for nodes on the saddles—into any of the three major brain states for each participant. The vectors that were not classified into any of the three major brain states were labelled as “other state”. This classification information would be used in the following EEG-triggered state-dependent TMS experiment.

#### 3.2.5. Offline EEG analysis: simulation of brain state dynamics

In the final part of the energy landscape analysis, we depicted and quantified the brain state dynamics by a random-walk simulation in the energy landscape ^19,23^. We simulated movement of the brain activity patterns using a Markov chain Monte Carlo method with the Metropolis-Hastings algorithm ^62,63^. Any brain activity pattern *V_i_* could move only to a neighbouring pattern *V_j_* with probability *P_ij_* = min[1,*e*^*E(V_i_)−E(V_j_*)]^. For each individual, we repeated this random walk 10^5^ steps with a randomly chosen initial pattern, which depicted the trajectory of activity patterns as a series of staying in and transitions between the three major brain states. The first 100 steps were discarded to minimise the influence of the initial condition.

Our previous work demonstrated a strong correlation between the percept duration and the length of return travel between Frontal state and Visual state via Intermediate state ^19^. Given this, we compared the behaviourally observed percept duration to the length of the Frontal–Intermediate–Visual travel that was calculated in the above random-walk simulation.

#### 3.2.6. Offline EEG analysis: temporal smoothing

In parallel with the random-walk simulation, we examined empirical brain state dynamics probing the binary neural vectors with seven elements, *V^t^*. For this purpose, we first categorised all *V^t^* into either of the three major brain states based on the classification information that was obtained in the above “Offline EEG analysis: structure of energy landscape” section. The vectors that were not classified into any of the three major brain states were labelled as “Other state”.

To reduce the effects of signal fluctuation, we then applied the following temporal smoothing to the time-series of the brain states. First, in a sliding window manner (window length = 10 msec), we calculated the appearance frequency for each brain state in the time window (Supplementary Fig. 3a), which was assigned to each centre time point in the window as the representative appearance frequency for each brain state (Supplementary Fig. 3b). Next, a Gaussian smoothing filter (FWHM = 10msec) was applied to the representative appearance frequency curves (Supplementary Fig. 3c). Based on the resultant appearance frequency values, we chose the most frequent brain state and assigned it as the brain state at the time point (Supplementary Fig. 3d). Note that this temporal smoothing eliminated the “other state”.

This empirical brain state dynamics would be used to evaluate the accuracy of the following online EEG analysis.

**Supplementary Fig. 3.**
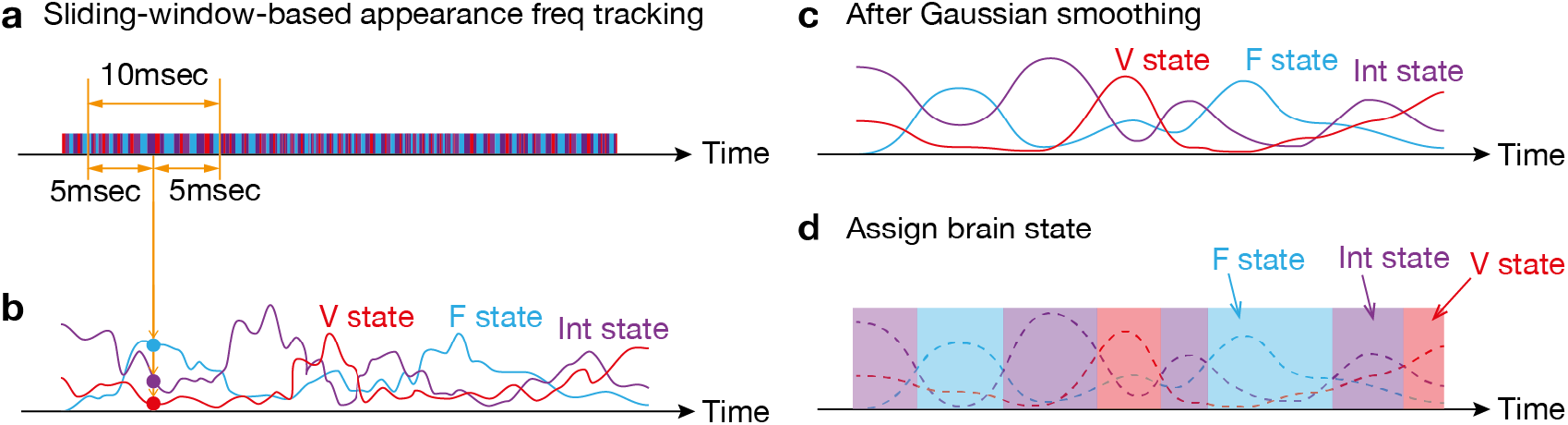
To reduce the effects of noise-oriented fluctuation in bran state dynamics, we applied two temporal smoothing procedures to the original brain state dynamics data. First, in a sliding window manner, we calculated three appearance frequency values for the three brain states (**panels a and b**). Then, Gaussian temporal smoothing was applied to the three appearance frequency curves (panel c). By comparing the smoothed curves, we identified which brain state was the most dominant at each timepoint (**panel d**).

#### 3.3.1. Online EEG analysis: preprocessing

The Online EEG analysis was conducted in all the state-dependent TMS experiments and its protocol was conceptually the same as that in the previous studies ^28–30,64^. The preprocessed EEG signals (<1.25kHz and 3kHz-downsampled) were input to the real-time target PC machine through a DAQ board (sampling rate, 2kHz), in which Simulink Real-Time model and xPC Target in Simulink (MathWorks, US) were running to analyse the EEG signals and trigger the TMS system via a TTL signal. We set the analysis model in the target PC through an Ethernet-connected host PC prior to the experiments for each participant.

First, using the “sum-of-difference” manner, we converted the EEG signals from 28 electrodes (4 electrodes/ROI × 7 ROIs) into seven Hjorth signals at each timepoint. Then, in a sliding-window manner (window length = 1000msec, i.e., 2000 time points), we extracted the gamma-band signals (30–80Hz) by applying a fast Fourier transformation (FFT) to EEG data in every time window (Supplementary Fig. 4a). The gamma band was chosen because (i) this EEG analysis aimed at reproducing our previous fMRI findings ^19^ and (ii) the EEG signals in the frequency window is considered to have temporal properties similar to those of fMRI signals ^57,59^. We then estimated the Hilbert envelope function for the gamma-band signals, whose amplitude would be used as neural signals ^57^(Supplementary Fig. 4b). To reduce the edge effects ^29,30^, we trimmed 100msec of the neural time-series data on both the edges of the time window (Supplementary Fig. 4c).

Using the remaining 800msec of the neural signals, we then conducted an autoregressive forward prediction based on Yule-Walker equation (order = 30)^29,30,64^ and estimated 280msec of neural signals that would come after the edge of the trimmed data (Supplementary Fig. 4d). That is, given the 100msec edge trimming, this calculation predicted the neural signal in a period between *T* = – 100msec and *T* = 180msec when we set *T* = 0 at the actual recording timing (i.e., the original terminal edge of the EEG data). The length of the forward prediction was set at 180msec because (i) one inhibitory TMS in this study took 150msec, (ii) the temporal smoothing required 5msec more data points for its sliding-window-based calculations (see Section 3.2.6), and (iii) we prepared 25-msec buffer for the following signal processing to realise a nearly simultaneous EEG-triggered TMS system^64^.

We obtained such 280-msec time-series signal for each ROI and used them in the following analysis.

**Supplementary Fig. 4.**
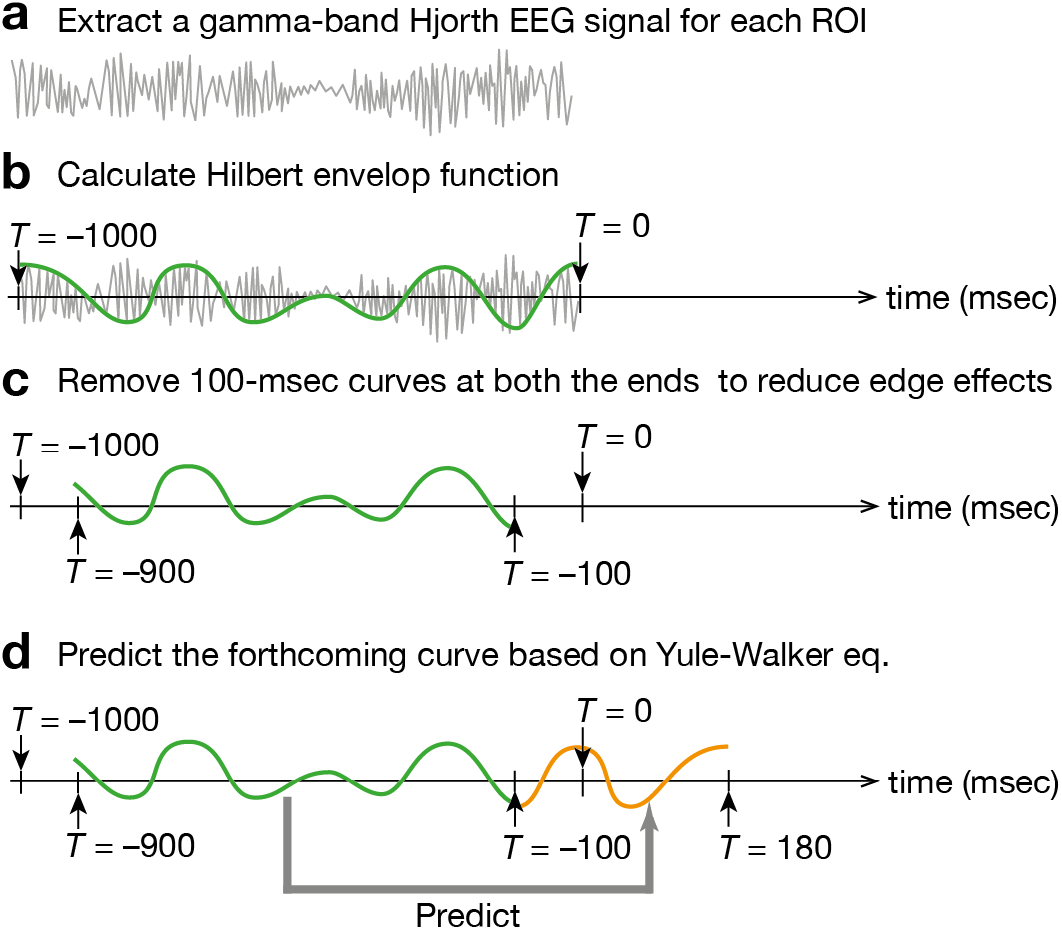
For the online preprocessing of EEG data, we first calculated a Hjorth EEG signal from each ROI data and then extracted a gamma-band signal in a sliding-window manner (**panel a**). Next, we fitted the Hilbert envelop function (**panel b**) and removed both ends of the 1000-msec time window to reduce the edge effects (**panel c**). Finally, based on the remaining 800-msec Hilbert envelop function, we made the prediction of the forthcoming 280-msec signal (**panel d**). Such 280-msec signals would be used for brain-state tracking.

#### 3.3.2. Online EEG analysis: binarisation

We then conducted online binarisation of the neural signals during the predicted time period. The binarisation threshold was calculated for each ROI based on the EEG data obtained in a control run that was conducted right before every TMS experiment. Technically, we applied the above-stated “Online EEG analysis: preprocessing” procedure to the EEG data during the control run and calculated the average of the Hilbert envelop amplitude for each ROI in each participant.

#### 3.3.3. Online EEG analysis: brain-state-/history-dependent TMS

These preprocessing and binarisation processes yielded a binary neural vector with seven elements (+1 or −1) at each time point in the 280-msec period (from *T* = –100 to *T* = 180). We then categorised the binary neural vectors into either of the three major brain states based on the classification information that was obtained in the offline energy landscape analysis in the control experiment. The binary neural vectors that were not categorised into any of the three major brain states were labelled as “Other state”.

Next, to reduce the effects of signal fluctuation, we applied the same temporal smoothing protocol to the resultant brain state vector as that used in the “Offline EEG analysis: temporal smoothing”. As a result of these analyses and smoothing procedures, we obtained a series of representative brain states in a period between *T* = −95msec and *T* = 175msec. Note that this temporal smoothing eliminated the “Other state”.

The state-dependent TMS was performed based on the brain states that were predicted to appear in the forthcoming period between *T* = 25msec and *T* = 175msec. The TMS was administered only when the target brain state was dominant in the period (>90% of the brain states). For example, for the Frontal-state-dependent TMS, only if Frontal state was predicted to dominantly appear in more than 90% of the time period, the real-time analysis machine sent a TTL trigger signal to the TMS device 21msec after the EEG signals were input to the analysis machine.

This 21msec buffer for the online EEG analysis was chosen because the EEG data took ~3msec to reach the analysis machine and the TTL signal from the analysis machine took ~1msec to trigger the TMS. Given these latencies, the TMS was supposed to start almost 25msec after the EEG recording (*T* = 25msec; Supplementary Fig. 5).

In addition, the 21msec buffer was sufficiently long for the online EEG analysis. By connecting the TTL signal back to the DAQ board of the real-time analysis machine, we evaluated the signal processing delay using the EEG data collected in the Control experiment I and confirmed that the TTL signal reached back to the analysis machine 21.3±0.02msec (mean±s.d., 21.1msec–21.5msec) after the EEG signals were input into the machine.

**Supplementary Fig. 5.**
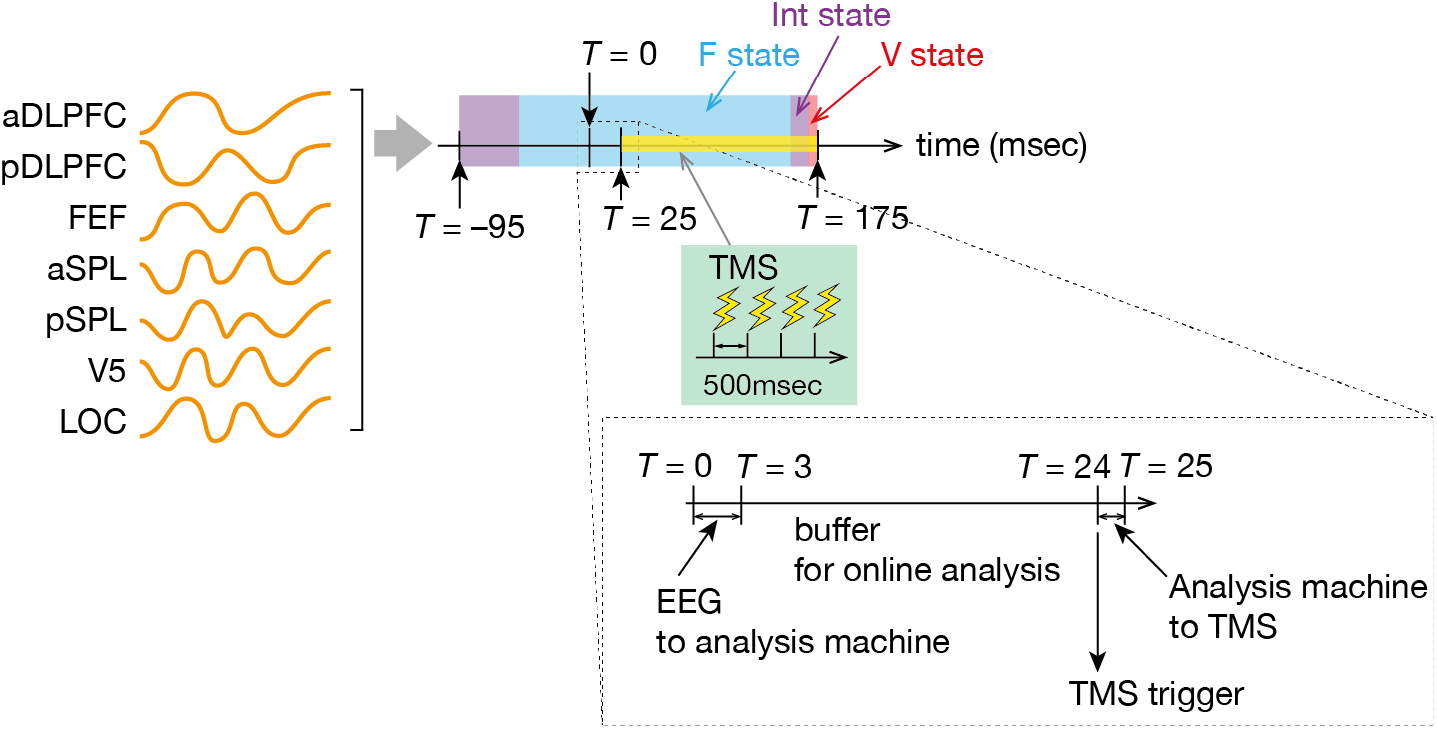
After the online preprocessing, we obtained seven time-series data for the seven ROIs (**orange curves in the left**). We put the seven ROI signals into binarisation, brain state categorisation and temporal smoothing procedures, which resulted in brain state dynamics in the 270-msec time period (*T* = −95 to *T* = 175 msec). When one brain state dominated the time window, the online analysis machine sent a trigger at *T* = 24msec after a 21-msec buffer period (**a dashed box in the bottom**). A burst of the inhibitory TMS consisted of three TMS stimulations with 500-msec intervals and thus took 150msec (**green box in the centre**).

For the neural-history-specific TMS, we looked into both the forthcoming period (*T* = 25msec to *T* = 175msec) and the preceding period (*T* = –95msec to *T* = 25msec). The TMS was administered only when both the two periods dominantly showed the target brain states. That is, for example, Post-Frontal Intermediate-specific TMS was administered only when Intermediate state was dominant in the forthcoming period (>90% of the brain states) and Frontal state was dominant in the neighbouring time window.

Note that, for both the brain-state-/history-specific stimulation, once a TMS was administered, we did not conduct the next stimulation until at least 9 sec passed. Such an interval gave the current brain-state-dependent TMS system a sufficient length of clean EEG data before the next TMS.

### 4.1. Experiment design and statistics

In the above sections, we stated all the essential device setups and analysis procedures. In the following sections, we elaborated on the actual designs of the experiments using such devices and analyses protocols.

#### 4.2.1. Control experiment

The control experiment consisted of two parts: EEG part and EEG/TMS part. The Control Experiment I was conducted to (i) identify the brain state dynamics in the offline EEG analysis and (ii) validate the locations of the EEG electrodes, whereas the Control Experiment II was performed to (iii) verify the accuracy of the online EEG analysis and brain-state-dependent TMS system. Both the parts employed the same 65 individuals and were performed with at least two-day intervals.

#### 4.2.2. Control Experiment I: EEG part (day 1)

In the Control Experiment I, we collected EEG signals while the participants were conducting the test of bistable visual perception (1.5min/run × 10 runs). The participants started this EEG recording sessions after sufficient training of the test.

For the aim (i), we applied the offline energy landscape analysis to the EEG data and examined whether the brain dynamics seen in our previous fMRI study ^19^ were qualitatively reproduced in the current EEG experiment. This analysis was conducted for the aim (ii) as well: if we successfully confirmed the reproducibility, such observations would provide face validation to the locations of the EEG electrodes. The details of this offline energy landscape were stated above (see “Offline EEG analysis” sections).

#### 4.2.3. Control Experiment II: EEG/TMS part (day 2)

In the Control Experiment II, the participants underwent test of bistable visual perception (1.5min/run × 11 runs) with the brain-state-dependent TMS system, which was almost the same as stated in the sections (see “Device setup: TMS” and “Online EEG analysis”) except for the locations of the TMS coil and an EEG electrode. We placed one of the 32 TMS-compatible Ag/AgCl ring electrodes—which was located on A1 in the original setting—on a wooden table that was set remotely from the participants. The TMS coil was placed over the electrode. Using the signal from this electrode, we measured when each TMS stimulation was conducted without causing significant artefacts on the EEG data from the participants.

The scalp EEG data in the first run were used to calculate the threshold value for binarisation. Therefore, we did not apply TMS in the run. In the rest ten runs, the TMS was performed in the following five different conditions: Frontal-/Intermediate-/Visual-state-dependent and Post-Frontal-/Post-Visual-Intermediate-state-dependent conditions. Each temporal condition was tested in two runs.

In the analysis, we performed both the offline and online analyses using the same EEG data. As stated above, the online EEG analysis required (i) the classification information to determine which major brain state would be assigned to each neural activity pattern and (ii) the threshold to binarise the neural signals. The requirement (i) was met by employing the results of the first half of the control experiment. For the requirement (ii), the data during in the first run were used: we conducted the preprocessing part of the online EEG analysis, extracted a Hilbert envelop curve and calculated the mean value of the envelope amplitude for each ROI in each participant.

Using the classification information and binarisation threshold, we performed the online analysis with the EEG data recorded during the rest of the runs (10 runs). This online EEG analysis was the same as that described above “Online EEG analysis” sections, and we obtained a time-series vector representing brain state dynamics.

For the sake of comparison, the offline analysis also used the EEG data recorded in the 2^nd^-11^th^ run and estimated the brain-state dynamics. We applied all the above-mentioned “Offline EEG analysis” procedures to the data except for the last random-walk simulation part. This offline analysis provided us with information about which major brain state should be assigned to each neural activity pattern.

Based on the classification information, we then labelled the preprocessed EEG time-series data, which resulted in a time-series vector of brain state dynamics.

In sum, these two types of EEG analysis gave us two time-series vectors representing the brain state dynamics for each participant. We then compared the two vectors and estimated how accurately the online analysis-based time-series vector predicted one that was based on the offline analysis. Technically, we counted timepoints whose brain states in the online analysis-based vector were the same as those in the offline analysis-based vector. We repeated such counting for each of the three major brain states in each participant and evaluated the accuracy.

In parallel, we examined the temporal accuracy of the TMS. The brain-state-dependent TMS system was designed to administer a TMS train 25msec after a specific brain state was detected in the EEG signals. Using the signal from the EEG electrode placed on the wooden table with the TMS coil over it, we measured the actual latency from the brain-state detection to the TMS stimulation and calculated the difference between the empirical latency and the presumed time-lag (i.e., 25msec). We repeated this estimation for all the five temporal conditions in each participant.

#### 4.3.1. Main experiment: design

In the main experiment, we examined causal behavioural effects of state-/state-history-dependent TMS on the median percept duration in a test of SFM-induced bistable visual perception. The participants underwent TMS over three different brain sites (aDLPFC, pDLPFC and FEF) in five different neural timings (F−. V−, Int−, Post-F Int- and Post-V Int-state-dependent TMS). In addition, we conducted an experiment using sham stimulation, in which mock TMS was applied over one of the three PFC sites. The choice of the target site in the sham condition was randomised and balanced across the participants.

These experiments were conducted on four different days with more than one-week intervals. In each day, the participants received TMS over the same brain sites in the five different brain-state conditions (Supplementary Fig. 6): they underwent six runs of bistable visual perception tests for each condition, between which they had one-min rest and one control run without TMS. The order of the brain-state conditions was randomised and balanced across the participants. Before and after the TMS conditions, the participants also performed longer control runs.

**Supplementary Fig. 6.**
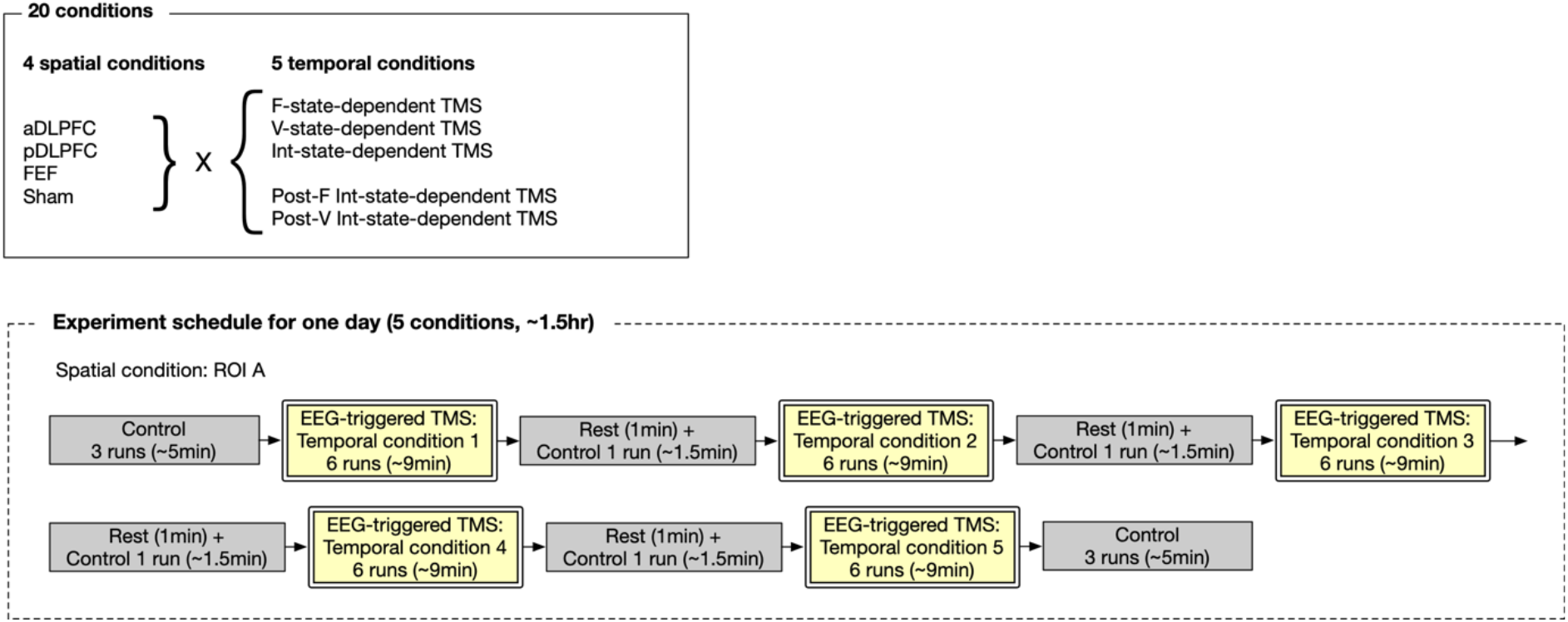
Design of the main experiment. For each participant, the experiment was conducted over four different days with >one-week intervals. On each day, they received TMS/sham over one of the three PFC days in five different timings. The stimulation site for the sham condition was randomly chosen from the three PFC areas and balanced across the participants.

#### 4.3.2. Main experiment: behavioural and neural analysis

Behaviourally, we calculated the median percept duration for each TMS condition and control sessions. The behavioural effects of the TMS were evaluated as the ratio of the median percept duration in the TMS conditions to that in the control runs conducted at the beginning of the entire main experiment.

Neurobiologically, we examined TMS’s effects on the brain state dynamics. Using the results of the online EEG analysis, we evaluated the dwelling time lengths for the three major brain states and transition frequencies between them after each TMS administration. To reduce the artefacts induced by the TMS, we discarded EEG signals recorded between the beginning of the TMS (*T* = 25msec) and 100msec after the end of the TMS (*T* = 175msec + 100msec = 275msec) and analysed the data in the first 4-sec time window in the remaining data (i.e., the period from *T* = 275msec to *T* = 4275msec).

#### 4.3.3. Main experiment: numerical simulation of TMS effects on energy landscape

To investigate the neurobiological changes in the brain state dynamics, we also numerically examined how the inhibitory TMS affected the structures of the individual energy landscape, such as the height of the energy barrier.

This simulation was conducted based on the *H_i_* and *J_ij_* that were obtained in the offline energy landscape analysis using the EEG data recorded during the control experiment. If ROI_*i*_ was the target site of the inhibitory TMS, we removed all the binary neural vectors *V^t^* whose element *i*, *σ_i_*, was set at +1 (i.e., active). Then, using the same *H_i_* and *J_ij_*, we built the disconnectivity graph (see “Offline EEG analysis: disconnectivity graph in energy landscape analysis”) and estimated the structural properties of the energy landscape (see “Offline EEG analysis: structure of energy landscape”). When the local minimum that represented any of the major brain states was removed in this simulation, we alternatively used the brain state that included the neighbouring neural vector as the major brain state. The neighbouring neural vector was defined as a vector that was different from the vector for the local minimum only in one element. In such numerical simulation, all the three major brain states were preserved whichever PFC site was inhibited.

#### 4.3.4. Main experiment: comparison between the numerical simulation, neural effects and behavioural causality

We compared the results of the numerical simulation, changes in the brain state dynamics and causal behavioural effects. First, we built hypotheses about the changes in the brain state dynamics based on the numerical simulation (Fig. 2h) and tested them by comparing the results of the numerical simulation with the effects on the brain state dynamics. After confirming the validity of the hypotheses, we examined the relationship between the energy landscape changes, effects on the brain state dynamics and causal behavioural responses using the mediation analysis.

### 4.4. Replication experiment

We examined the replicability of the findings of the main experiments by repeating the main experiment with focusing on the seven conditions that yielded significant behavioural effects. This experiment employed a small but independent cohort (originally *N*=15; *N*=14 after one participant dropped out).

### 4.5. Validation experiment with excitatory TMS

We also examined the validity of the main findings in experiments using the excitatory TMS (see details on the TMS protocol in “Device setup: TMS” section). As shown in our previous work ^35,36^, the excitatory TMS was expected to induce behavioural and neural effects opposite to those yielded by the inhibitory TMS. To test this, we repeated the main experiment using the excitatory TMS with an independent cohort (*N*=15). Like the replication experiment, we focused on the seven TMS conditions that induced significant behavioural changes in the main experiment.

### 4.6. Conventional quadripulse TMS experiment

For comparison, we examined the behavioural effects of conventional 30-min inhibitory quadripulse TMS (QPS) by employing the 14 individuals who completed the replication experiment at least one month before.

In this experiment, we adopted the inhibitory QPS protocols that were used in our previous studies ^35,36^. This TMS consisted of 360 consecutive bursts with 5-sec intervals (i.e., 30min), and each burst comprised four monophasic 20Hz TMS whose intensity was set at 90% of AMT. We conducted this QPS over either of the three PFC regions using the same TMS device as used in the main experiment. In addition, we performed a sham condition; thus, the participants came to the lab four times with more-than-one-week interval. The order of the TMS conditions was randomised and balanced across the participants.

In each day, the participants first completed the six control runs of bistable visual perception tests and then underwent the 30min TMS session over one of the three PFC sites. In the sham condition, the TMS coil was placed over one of the three PFC areas, which was randomly chosen and balanced across the participants. During the TMS session, they were asked to rest with their eyes open. Ten minutes after the end of the TMS, they took the six runs of the bistable visual perception tests.

### 4.7. Longitudinal experiment

We investigated accumulative effects of the brain-sate-/history-specific TMS by a longitudinal TMS experiment that employed the 63 participants who completed the main, replication or validation experiment.

Like the replication and validation experiments, we focused on the seven TMS conditions that induced the significant behavioural changes. In addition, we added three sham conditions, in which the TMS was administered over one of the three PFC regions at random timings. In total, the ten conditions were examined.

The 63 participants were randomly assigned to two of the ten conditions: each non-sham condition had 13 individuals, whereas two sham condition had 12 participants and the other sham condition was examined with 11 individuals.

The accumulative effect of each TMS/sham condition was evaluated over nine weeks. In the first eight weeks, the participants underwent weekly TMS/sham experiments. In each day, the participants underwent a TMS session (six runs of bistable visual perception tests) and two control session (three runs) before and after the TMS session. The details of the TMS protocol were the same as those for the main experiment. The brain site for the sham condition was randomly chosen from the three PFC areas for each participant for each session. The target brain sites were not changed during one period.

For each day, we analysed the behavioural effects of the state-/state-history-dependent TMS by calculating proportional changes in the median percept durations within each day. We also tracked the behavioural effects over the two-month experiment and examined whether the magnitude of the behavioural response at each day was significantly different from the first day (week 0).

In the last week (week 9), the participants underwent the control sessions only. Such baseline responses were used to evaluate whether the two-month weekly TMS affected the baseline perceptual stability.

## 5. Statistics

In the case of multiple comparisons, basically, we conducted a two-way ANOVA (participant × condition) and post-hoc *t*-tests with Bonferroni correction.

## Acknowledgements

This work was supported by Grant-in-aid for Research Activity from Japan Society for Promotion of Sciences, The University of Tokyo Excellent Young Researcher Project, Fukuhara Foundation, Yamaha Motor Foundation for Sports, Astellas Foundation for Research on Metabolic Disorders, Showa University Medical Institute of Developmental Disabilities Research and JST Moonshot R&D Program (JPMJMS2021). Accessibility to some experimental facilities used here was realised through financial support from Yasushi Miyashita’s team in RIKEN CBS, Japan. The author appreciates stimulating hints and sharp insights of Professor Geraint Rees (UCL, UK) in the initial phase of this research.

## 6. Data and code availability

The behavioural data are deposited in Dryrad (https://doi.org/10.5061/dryad.8931zcrqn) and the codes for the energy landscape analysis has been shared as a supplementary information of our previous work ^24^.

## References

1. Sterzer, P., Kleinschmidt, A. & Rees, G. The neural bases of multistable perception. Trends Cogn Sci 13, 310–318 (2009).

2. Brascamp, J., Sterzer, P., Blake, R. & Knapen, T. Multistable Perception and the Role of Frontoparietal Cortex in Perceptual Inference. Annu Rev Psychol 69, 1–27 (2017).

3. Lumer, E. D., Friston, K. J. & Rees, G. Neural Correlates of Perceptual Rivalry in the Human Brain. Science 280, 1930–1934 (1998).

4. Kleinschmidt, A., Bchel, C., Zeki, S. & Frackowiak, R. S. J. Human brain activity during spontaneously reversing perception of ambiguous figures. Proc Royal Soc Lond Ser B Biological Sci 265, 2427–2433 (1998).

5. Sterzer, P., Russ, M. O., Preibisch, C. & Kleinschmidt, A. Neural Correlates of Spontaneous Direction Reversals in Ambiguous Apparent Visual Motion. Neuroimage 15, 908–916 (2002).

6. Wang, M., Arteaga, D. & He, B. J. Brain mechanisms for simple perception and bistable perception. Proc National Acad Sci 110, E3350–E3359 (2013).

7. Weilnhammer, V. A., Ludwig, K., Hesselmann, G. & Sterzer, P. Frontoparietal Cortex Mediates Perceptual Transitions in Bistable Perception. J Neurosci 33, 16009–16015 (2013).

8. Panagiotaropoulos, T. I., Deco, G., Kapoor, V. & Logothetis, N. K. Neuronal Discharges and Gamma Oscillations Explicitly Reflect Visual Consciousness in the Lateral Prefrontal Cortex. Neuron 74, 924–935 (2012).

9. Panagiotaropoulos, T. I., Dwarakanath, A. & Kapoor, V. Prefrontal Cortex and Consciousness: Beware of the Signals. Trends Cogn Sci 24, 343–344 (2020).

10. Lumer, E. D. & Rees, G. Covariation of activity in visual and prefrontal cortex associated with subjective visual perception. Proc National Acad Sci 96, 1669–1673 (1999).

11. Hohwy, J., Roepstorff, A. & Friston, K. Predictive coding explains binocular rivalry: An epistemological review. Cognition 108, 687–701 (2008).

12. Weilnhammer, V., Stuke, H., Hesselmann, G., Sterzer, P. & Schmack, K. A predictive coding account of bistable perception - a model-based fMRI study. Plos Comput Biol 13, e1005536 (2017).

13. Graaf, T. A. de, Jong, M. C. de, Goebel, R., Ee, R. van & Sack, A. T. On the Functional Relevance of Frontal Cortex for Passive and Voluntarily Controlled Bistable Vision. Cereb Cortex 21, 2322–2331 (2011).

14. Harrison, S. A. & Tong, F. Decoding reveals the contents of visual working memory in early visual areas. Nature 458, 632–635 (2009).

15. Brascamp, J., Blake, R. & Knapen, T. Negligible fronto-parietal BOLD activity accompanying unreportable switches in bistable perception. Nat Neurosci 18, 1672–1678 (2015).

16. Knapen, T., Brascamp, J., Pearson, J., Ee, R. van & Blake, R. The Role of Frontal and Parietal Brain Areas in Bistable Perception. J Neurosci 31, 10293–10301 (2011).

17. Frässle, S., Sommer, J., Jansen, A., Naber, M. & Einhäuser, W. Binocular Rivalry: Frontal Activity Relates to Introspection and Action But Not to Perception. J Neurosci 34, 1738–1747 (2014).

18. Block, N. Finessing the Bored Monkey Problem. Trends Cogn Sci 24, 167–168 (2020).

19. Watanabe, T., Masuda, N., Megumi, F., Kanai, R. & Rees, G. Energy landscape and dynamics of brain activity during human bistable perception. Nat Commun 5, 4765 (2014).

20. Silvanto, J., Muggleton, N. & Walsh, V. State-dependency in brain stimulation studies of perception and cognition. Trends Cogn Sci 12, 447–454 (2008).

21. Bergmann, T. O. Brain State-Dependent Brain Stimulation. Front Psychol 9, 2108 (2018).

22. Zrenner, C., Belardinelli, P., Müller-Dahlhaus, F. & Ziemann, U. Closed-Loop Neuroscience and Non-Invasive Brain Stimulation: A Tale of Two Loops. Front Cell Neurosci 10, 92 (2016).

23. Watanabe, T. & Rees, G. Brain network dynamics in high-functioning individuals with autism. Nat Commun 8, 16048 (2017).

24. Ezaki, T., Watanabe, T., Ohzeki, M. & Masuda, N. Energy landscape analysis of neuroimaging data. Philosophical Transactions Royal Soc Math Phys Eng Sci 375, 20160287 (2017).

25. Kang, J., Pae, C. & Park, H.-J. Energy landscape analysis of the subcortical brain network unravels system properties beneath resting state dynamics. Neuroimage 149, 153–164 (2017).

26. Gu, S. et al. The Energy Landscape of Neurophysiological Activity Implicit in Brain Network Structure. Sci Rep-uk 8, 2507 (2018).

27. Bergmann, T. O., Karabanov, A., Hartwigsen, G., Thielscher, A. & Siebner, H. R. Combining non-invasive transcranial brain stimulation with neuroimaging and electrophysiology: Current approaches and future perspectives. Neuroimage 140, 4–19 (2016).

28. Schaworonkow, N., Triesch, J., Ziemann, U. & Zrenner, C. EEG-triggered TMS reveals stronger brain state-dependent modulation of motor evoked potentials at weaker stimulation intensities. Brain Stimul 12, 110–118 (2019).

29. Stefanou, M.-I., Desideri, D., Belardinelli, P., Zrenner, C. & Ziemann, U. Phase Synchronicity of μ-Rhythm Determines Efficacy of Interhemispheric Communication Between Human Motor Cortices. J Neurosci 38, 10525–10534 (2018).

30. Zrenner, C., Desideri, D., Belardinelli, P. & Ziemann, U. Real-time EEG-defined excitability states determine efficacy of TMS-induced plasticity in human motor cortex. Brain Stimul 11, 374–389 (2018).

31. Miller, S. M. et al. Interhemispheric switching mediates perceptual rivalry. Curr Biol 10, 383–392 (2000).

32. Pettigrew, J. D. & Miller, S. M. A sticky interhemispheric switch in bipolar disorder? Proc Royal Soc Lond Ser B Biological Sci 265, 2141–2148 (1998).

33. Haynes, J.-D., Deichmann, R. & Rees, G. Eye-specific effects of binocular rivalry in the human lateral geniculate nucleus. Nature 438, 496–499 (2005).

34. Hamada, M. et al. Quadro-pulse stimulation is more effective than paired-pulse stimulation for plasticity induction of the human motor cortex. Clin Neurophysiol 118, 2672–2682 (2007).

35. Watanabe, T. et al. Bidirectional effects on interhemispheric resting-state functional connectivity induced by excitatory and inhibitory repetitive transcranial magnetic stimulation. Hum Brain Mapp 35, 1896–1905 (2014).

36. Watanabe, T. et al. Effects of rTMS of Pre-Supplementary Motor Area on Fronto Basal Ganglia Network Activity during Stop-Signal Task. J Neurosci 35, 4813–4823 (2015).

37. Hamada, M. et al. Primary motor cortical metaplasticity induced by priming over the supplementary motor area. J Physiology 587, 4845–62 (2009).

38. Karabanov, A., Thielscher, A. & Siebner, H. R. Transcranial brain stimulation. Curr Opin Neurol 29, 397–404 (2016).

39. Cattaneo, L., Sandrini, M. & Schwarzbach, J. State-Dependent TMS Reveals a Hierarchical Representation of Observed Acts in the Temporal, Parietal, and Premotor Cortices. Cereb Cortex 20, 2252–2258 (2010).

40. Cattaneo, Z., Devlin, J. T., Salvini, F., Vecchi, T. & Silvanto, J. The causal role of category-specific neuronal representations in the left ventral premotor cortex (PMv) in semantic processing. Neuroimage 49, 2728–2734 (2010).

41. Ezzyat, Y. et al. Direct Brain Stimulation Modulates Encoding States and Memory Performance in Humans. Curr Biol 27, 1251–1258 (2017).

42. Scangos, K. W., Makhoul, G. S., Sugrue, L. P., Chang, E. F. & Krystal, A. D. State-dependent responses to intracranial brain stimulation in a patient with depression. Nat Med 27, 229–231 (2021).

43. Polanía, R., Nitsche, M. A. & Ruff, C. C. Studying and modifying brain function with non-invasive brain stimulation. Nat Neurosci 21, 174–187 (2018).

44. Mrachacz-Kersting, N. et al. Brain state–dependent stimulation boosts functional recovery following stroke. Ann Neurol 85, 84–95 (2019).

45. Ezaki, T., Sakaki, M., Watanabe, T. & Masuda, N. Age-related changes in the ease of dynamical transitions in human brain activity. Hum Brain Mapp 39, 2673–2688 (2018).

46. Watanabe, T., Lawson, R. P., Walldén, Y. S. E. & Rees, G. A neuroanatomical substrate linking perceptual stability to cognitive rigidity in autism. J Neurosci 39, 2831–18 (2019).

47. Wassermann, E. M. Risk and safety of repetitive transcranial magnetic stimulation: report and suggested guidelines from the International Workshop on the Safety of Repetitive Transcranial Magnetic Stimulation, June 5-7, 1996. Electroencephalogr Clin Neurophysiology Evoked Potentials Sect 108, 1–16 (1998).

48. Rossi, S., Hallett, M., Rossini, P. M., Pascual-Leone, A. & Group, T. S. of T. C. Safety, ethical considerations, and application guidelines for the use of transcranial magnetic stimulation in clinical practice and research. Clin Neurophysiol 120, 2008–2039 (2009).

49. Freeman, E. D., Sterzer, P. & Driver, J. fMRI correlates of subjective reversals in ambiguous structure-from-motion. J Vision 12, 35–35 (2012).

50. Kanai, R., Bahrami, B. & Rees, G. Human Parietal Cortex Structure Predicts Individual Differences in Perceptual Rivalry. Curr Biol 20, 1626–1630 (2010).

51. Kanai, R., Carmel, D., Bahrami, B. & Rees, G. Structural and functional fractionation of right superior parietal cortex in bistable perception. Curr Biol 21, R106–R107 (2011).

52. Carmel, D., Walsh, V., Lavie, N. & Rees, G. Right parietal TMS shortens dominance durations in binocular rivalry. Curr Biol 20, R799–R800 (2010).

53. Hjorth, B. EEG analysis based on time domain properties. Electroen Clin Neuro 29, 306–310 (1970).

54. Seeck, M. et al. The standardized EEG electrode array of the IFCN. Clin Neurophysiol 128, 2070–2077 (2017).

55. Shirota, Y. et al. Cerebellar dysfunction in progressive supranuclear palsy: a transcranial magnetic stimulation study. Movement Disord 25, 2413–9 (2010).

56. Hanajima, R. et al. Interhemispheric facilitation of the hand motor area in humans. J Physiology 531, 849–859 (2001).

57. Watanabe, T., Rees, G. & Masuda, N. Atypical intrinsic neural timescale in autism. Elife 8, e42256 (2019).

58. Delorme, A. & Makeig, S. EEGLAB: an open source toolbox for analysis of single-trial EEG dynamics including independent component analysis. J Neurosci Meth 134, 9–21 (2004).

59. Deligianni, F., Centeno, M., Carmichael, D. W. & Clayden, J. D. Relating resting-state fMRI and EEG whole-brain connectomes across frequency bands. Front Neurosci-switz 8, 258 (2014).

60. Jaynes. Information theory and statistical mechanics. (1957).

61. Watanabe, T. et al. A pairwise maximum entropy model accurately describes resting-state human brain networks. Nat Commun 4, 1370 (2013).

62. Massen, C. P. & Doye, J. P. K. Identifying communities within energy landscapes. Phys Rev E 71, 046101 (2005).

63. Girvan, M. & Newman, M. E. J. Community structure in social and biological networks. Proc National Acad Sci 99, 7821–7826 (2002).

64. Chen, L. L., Madhavan, R., Rapoport, B. I. & Anderson, W. S. Real-Time Brain Oscillation Detection and Phase-Locked Stimulation Using Autoregressive Spectral Estimation and Time-Series Forward Prediction. Ieee T Bio-med Eng 60, 753–762 (2013).

